# Structural Basis for DNA Replication and Uracil Repair in Phage A-Family DNA Polymerases

**DOI:** 10.1101/2025.03.26.645540

**Authors:** Sophia Missoury, Soizick Lucas-Staat, Rémi Sieskind, Marc Delarue

## Abstract

Replicative DNA polymerases (DNAP) play a critical role in genome duplication, ensuring the accurate transmission of genetic information in all kingdoms of life. This process is essential also for DNA-dependent viruses, including bacteriophages. In many phage genomes, a uracil-DNA glycosylase (UDG) is encoded in trans. In this paper, we identify a new subfamily of A-family DNAP in phages that are fused to an active (UDG) domain. Two members of this subfamily, *B. subtilis* phage SP-15 and YerA41 are known to be hypermodified on their thymidines. Here, we present cryo-EM structures at high resolution for two of its members in various functional and conformational states, from YerA41 and phiLo phages. The structures explain how these DNAPs can have an activity, distinct from copying genetic information, which reads dU bases ahead of the replication fork and creates abasic sites that are efficiently bypassed by the DNAP. Additionally, we report the co-occurrence of both a X-family DNAP and a DNA ligase in the corresponding phage genomes, and show that both enzymes are capable of repairing the abasic sites. The former makes a stable complex with the replicative DNAP, which thus appears as a platform for recruitment of the enzymatic activities necessary for the repair of dU bases during replication, that result from the incorporation of residual dUTP in the pool of nucleotides. Strikingly, the location of the UDG domain is the same as in the structure of Mpox virus replicative complex involving a B-family DNAP.

## Introduction

The introduction of chemical modifications to some DNA bases is a widespread strategy used by phages to prevent the processing and destruction of their genetic material by host endonucleases^1–9^. Hosts can develop defense mechanisms and recent studies have revealed a variety of strategies employed by phages to counter these bacterial defense mechanisms for their survival^3,5,10^. Notably, very few of these modifications affect the Watson-Crick base-pairing positions^11–14^. Instead, most of them occur on the DNA backbone or positions on the bases that do not interfere with base pairing^8^. However, such modifications cause significant challenges to sequencing technologies in metagenomics projects, rendering some genome unsequencable with standard methods and hinting at the possible existence of regions of “genetic dark matter”^15,16^.

To counteract these DNA modifications, bacteria have evolved specific glycosylases. For example, bacterial glycosylases can excise α-glycosyl-hydroxymethylcytosine (5-hmC) from T4 phage DNA, preventing infection^17^. Other phages such as T2, T6 and RB69 also contain glycosylated 5-hmC^18^. Uracil DNA glycosylases (UDG) are also present in viruses and phages: for instance UDG protein was recently reported in 10 cases out of 22 in metagenomics studies of Crass-like phages^19^, underscoring their critical role in the evolutionary arms race between bacteria and phages^20,21^. UDG cleaves deoxyuridine (dU) base when present into the genomic DNA and leaves an abasic site^22^. dU, a modified base, can result from the oxidative deamination of cytosine, leading to mutations if left unrepaired or it may arise from the misincorporation by the phage DNAP of dUTP, which can be present in significant amount in the host’s dNTP pool. This last situation is actually one of the main counter-measures of the host, namely an increase of dUTP concentration in the cell, leading to dU incorporation that will render the DNA phage again accessible to endonucleases. It is then essential for the phage to have a way to get rid of dU bases. The abasic site created by UDG is typically corrected through the base excision repair (BER) pathway, involving enzymes such as AP lyase, AP endonucleases, X-family DNA polymerase (DNAP X), and DNA ligase ^23–26^. Whether these activities are encoded by the phage itself or rely by the host - either as separate genes or as multifunctional enzymes - varies significantly across different systems.

Physical interactions between replicative B-family DNA polymerases (DNAPs) and uracil-DNA glycosylases (UDG) encoded *in trans* have been observed in DNA viruses that infect eukaryotic cells, such as Herpesviruses, Monkeypox virus, and African swine fever virus^27^. These viral UDGs interact directly with the replicative DNA polymerase and the single-stranded DNA template ahead of the replication process^15–17^, suggesting a specific link between DNA repair and replication. In A-family DNA polymerases a recent reclassification has identified a subfamily of DNAP as a distinct cluster, referred to as N4-like polymerases, which contains a fused UDG-domain within the DNA polymerases^31^. In this subfamily, the UDG-domain has lost the conserved catalytic amino acids. However, DNAPs from phages such as those of SPO1, SP-10 or Shanette are known to contain active fused-UDG domains, although they do not appear as a distinct cluster in the current CLANS classification of A-family DNAPs^31^. Here, the interactions between the UDG domain and DNAP itself are unknown.

In this study, we present a structural and functional characterization of two A-family DNA polymerases fused with active UDG domains, both of which contain three distinct substantial insertions. One DNAP originates from a phage that is known to be T-modified by glycosylation, while the other is from a thermophilic phage whose DNA modification is not yet characterized. Using cryogenic-electron microscopy, we solved the three-dimensional structure of these polymerases in different conformational states, providing detailed insights into their domain structural organization. This analysis revealed the role of these novel insertions, critical for enzymatic activity, as well as the position and the function of the UDG-domain. Genome analysis of these phages uncovered the presence of intrinsic DNA repair enzymes, including DNAP X and DNA ligase, highlighting a self-sufficient repair system. We confirmed their repair functions on newly synthesized dsDNA containing abasic sites. This supports the existence of an evolved mechanism that strategically integrates DNA replication and repair and enhances phage and virus genome stability in response to host defense systems, probably valid for both A- and B-family DNAPs.

## Results

### Additional domains in A-family DNA polymerases encoded in phages

Czernecki *et al*. have proposed a revised classification for A-family DNA polymerases, specifically identifying two distinct clusters, each fused with an additional N-terminal domain identified as uracil DNA glycosylases (UDGs)^31^. The first cluster belongs to the N4-like DNA polymerases, which have lost their catalytic residues in the UDG domain. In contrast, the second cluster consists of SPO1-like DNA polymerases, where the catalytic residues of the UDG domain are conserved, as seen in DNA polymerases from SPO1 and SP-10 (**Fig. 1a**)^31^. Since this classification was reported, a large DNA polymerase from the **YerA41** phage has been discovered, containing 1,306 amino acids and a N-terminal UDG-like domain^8^. The cloning and purification of this DNAP (identified by RNA-seq) was essential in the determination of the genome, that resisted classical amplification methods^16^. CLANS analysis shows that it does not cluster with either N4-like or SPO1-like sequences^32^. Using BLAST, we identified related DNAPs from phages such as **phiLo**, a presumably temperature-resistant phage since its host is *Thermus thermophilus*, as well as RP13, LiS04 and SP-15 (**Fig. 1a-b**). SP-15 is known for its high-level of DNA modification, involving the addition of a glucosyl-phosphoglucuronolactone to a 5- dihydroxypentyl-uracil (DHPU) nucleobase, predicted to append a total of 488 Da side chain to thymidine nucleotides^33^. The bridging moiety between uracil and the sugars is 4’-5’ dihydroxypentyl (DHP) in SP-15 phage. Genomic DNA is also highly modified in *Ralstonia* phage RP13, although the nature of its modifications has not been precisely identified, and the authors mention that the sequence of the genome also had to proceed through RNA-seq sequencing because commercial DNA polymerases could not be used to amplify the genome, exactly as in YerA41^15^. RP13 genome contains a gene involved in thymidine modification, aGPT-PlpaseI (g120), alpha-glutamyl-putrescinyl thymidine pyrophosphorylase ^34^, which is also present in EP_H11 and EcoM_IME92. However, the often associated 5-hydroxymethyl-uridine DNA kinase (5HMUDK) could not be detected ^7,35^. In YerA41 several experimental evidence indicate that its genome contains a majority of glycosylated dT bases, in particular mass spectrometry detects a modification of thymidine with an additional mass of 1044 Da with a probable presence of up to five sugars^8^. The mass of the spacer between the 5-position of the pyrimidine ring and the sugars was measured to be 104 Da, 16 Da more than DHP in SP-15 ^36^. Overall, the clustering of SP-15 DNAP with DNAPs from YerA41 and RP13 phages and others (**Fig. 1a-b**), strongly suggests that these bacteriophages exhibit similar modifications to their DNA involving glycosylation of T and a longer linker than just the 5-hydroxy-methyl group (5hm)^33,37^.

**Figure 1.**
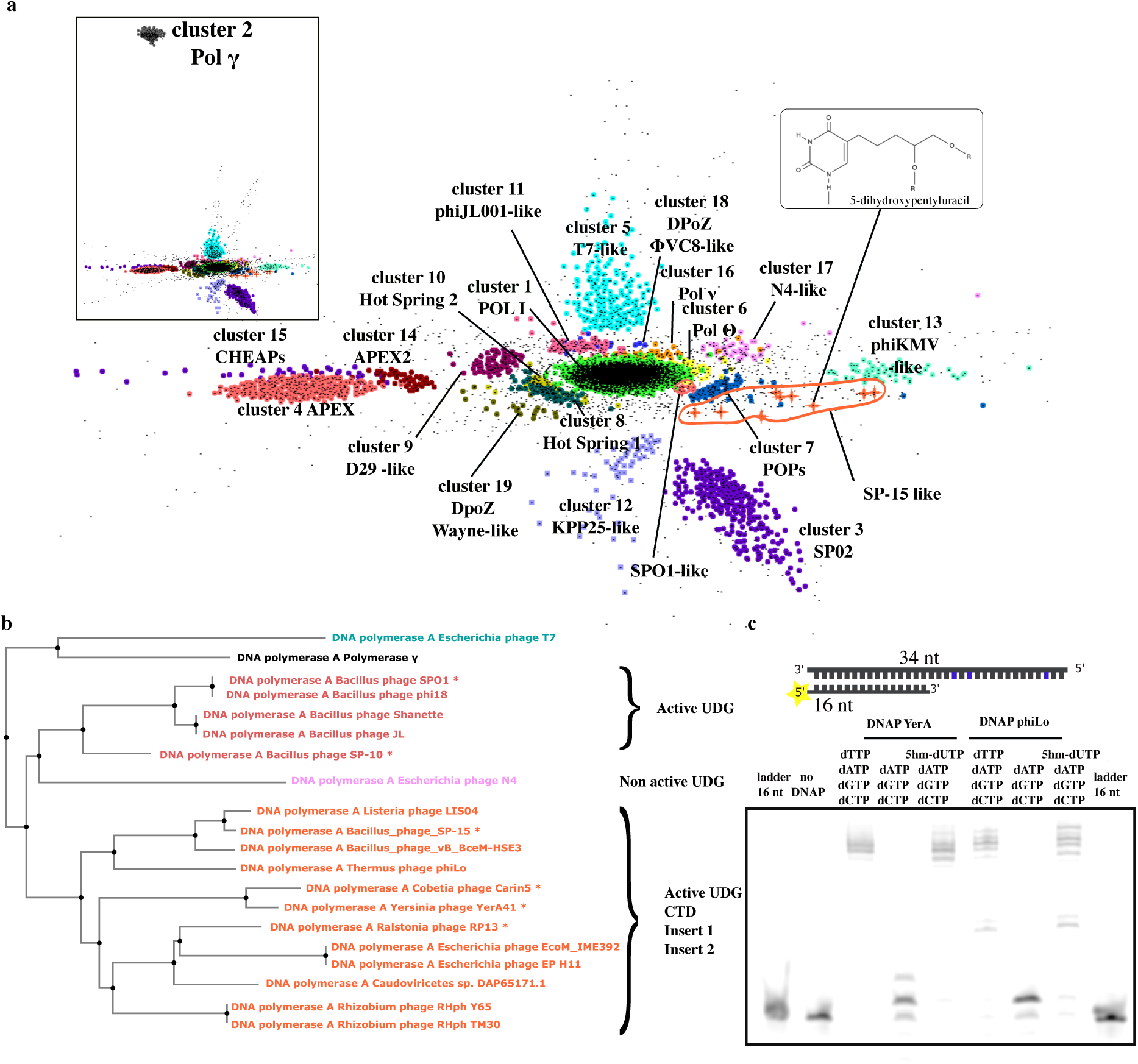
Positioning of the SP15-like set of DNAPs among all A-family DNAPs. **a.** Re-clustering of DNA polA sequences as described in Czernecki *et al*. (2023), this time incorporating additional DNAPs sequences from YerA41 and SP15-like phages identified through BlastP. The 3D distribution of the PolA members was generated using CLANS^32^, based on Blast pairwise scores between polymerase sequences converted into binary distances. The 19 major clusters of PolA sequences are color-coded as in the referenced paper. The SPO1-like cluster (too small to be identified in the original study) and SP15-like sub-family are highlighted in orange. Upper-left insert: cluster #2 (Pol *γ*) is widely separated from all the other clusters. Upper-right insert: The modified base 5-dihydroxypentyluracil observed in SP-15 genome. **b.** Distance tree generated from Blast searches showing the relationships between the SP15-like members and reference DNA polymerases, including bacteriophage T7 DNAP (cyan), mitochondrial pol *γ* (black), bacteriophage N4 pol (pink), and SPO1-like DNAPs with fused active UDG domains (red). SP15-like members are distinct from the reference sequences and colored in orange; they are further annotated as containing 3 domain/insertion(s) (CTD, Insert 1 and Insert 2). Organisms known to contain modified bases into their genome are annotated with a star. c. The primer extension activity of DNAP YerA41 and DNAP phiLo was evaluated using a 5’-FAM-labeled primer DNA substrate and template DNA strand containing dA bases in blue. The denaturing gel was scanned for FAM fluorescence, probing the progression of DNA synthesis in different conditions, as indicated on top of the lanes.

T-modifications have been studied extensively in several phages or viruses^7,35^ and they primarily involve the replacement of dTTP in the pool of dNTPs by 5hm-dUTP, synthesized through the successive action of a dCTP deaminase, a dUTP diphosphorylase and a thymidine synthase homologue. The incorporation of 5hm-dUTP by the DNA polymerase serves as a precursor for subsequent branching of a linker to the hydroxyl group, potentially followed by its glycosylation. In DNAPs from YerA41 and phiLo, a primer extension assay was performed in presence of 5hm-dUTPs instead of dTTPs and it confirmed the ability of SP15-like DNAPs to incorporate the 5hm-dUTP precursor (**Fig. 1c**). A multiple sequence alignment (MSA) of the members of this newly identified cluster of DNAPs (**Suppl. Fig. 1**) revealed conserved catalytic residues in the UDG-domain, annotated as motif UDG-A and motif UDG-B in yellow in **Suppl. Fig. 1**, suggesting potential enzymatic activity for this domain, akin to SPO1 and SP-10 DNA polymerases. Additionally, theses sequences contain an extended region at the C-terminal domain (CTD), consisting of around 80 amino acids, which is absent in both SPO1- like or N4-like polymerases (**Fig. 2a, Suppl. Fig. 1, Suppl. Fig. 2a-c)**. Given the existence of this distinct region and their divergence from N4-like and SPO1-like sequences, we propose that these DNAPs represent a new subfamily, which we designate as SP15-like DNAPs (**Suppl. Fig. 2a-b**).

**Figure 2.**
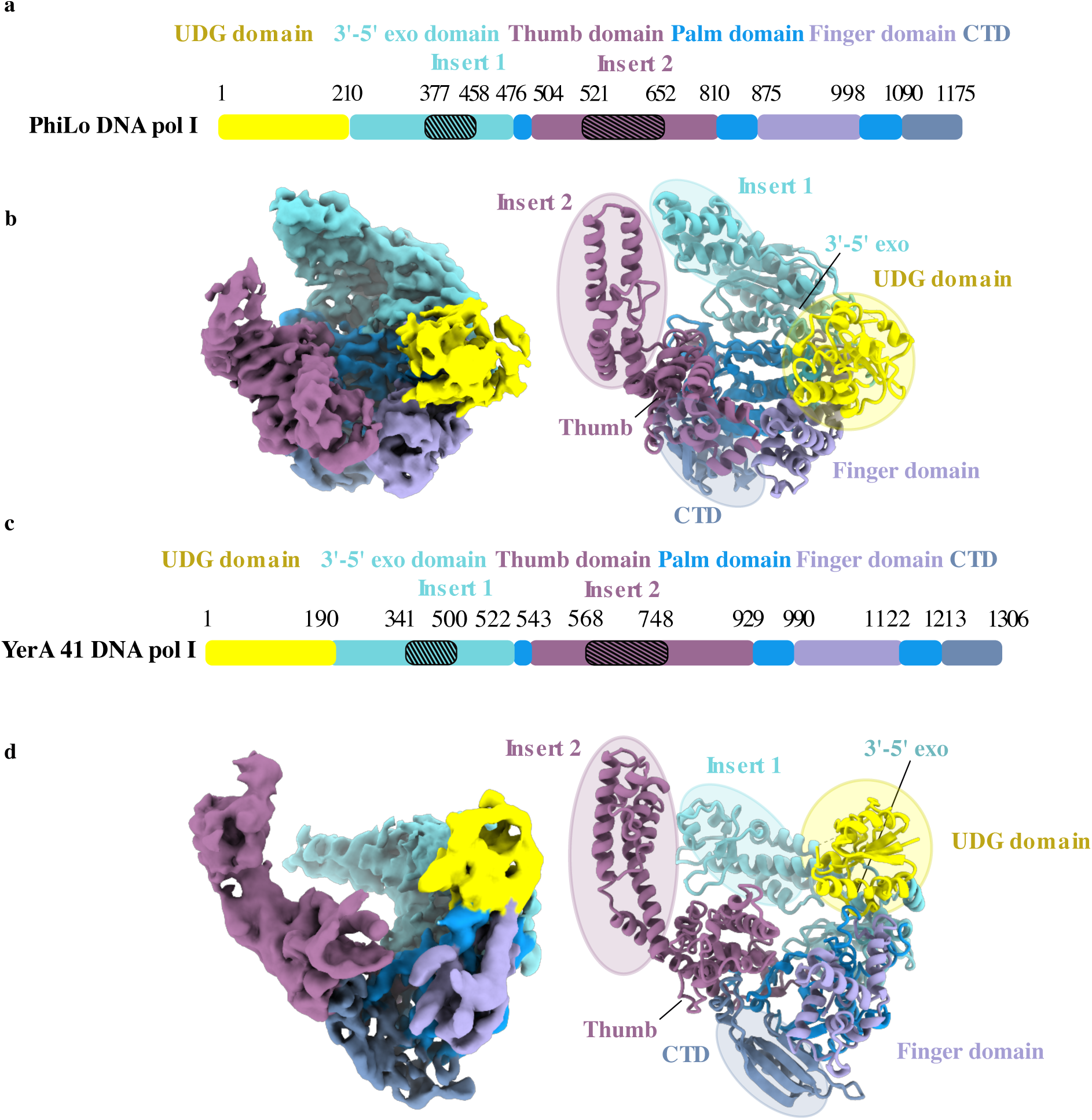
Additional domains in SP15-like A-family DNA polymerases. **a.** Schematic representation of the domain organization of phiLo DNA polymerase. The domains—UDG, 3′-5′ exonuclease (3′-5′ exo), thumb, palm, finger, and C-terminal (CTD)—are color-coded in yellow, cyan, light blue, blue, purple, and dark blue, respectively. The large insertions are hashed. The residue numbers at the borders of domains are indicated. **b.** Cryo-EM density map of phiLo DNAP in the apo conformation, with a ribbon model in the same orientation on the right, with domains color-coded as in (a) and corresponding domain annotations. **c.** Domain organization of YerA41 DNAP, as in (a). **d.** Cryo-EM map of YerA41 DNAP in the apo conformation, with a ribbon model on the right in a comparable orientation for reference. The UDG domain, Insert 1 of the 3′-5′ exonuclease domain, and Insert 2 of the thumb domain are labeled. The UDG domain (yellow) is positioned above the finger domain (purple). Inserts 1 and 2, which extend near the DNA binding site entry, are conserved in both DNA polymerases homologs.

In the following we present their three-dimensional structure, focusing on YerA41 DNAP and phiLo DNAP, which exhibit only 28% sequence identity, reflecting the high diversity commonly observed among phage DNA polymerases. We note that structure prediction of these DNA polymerases using AlphaFold^38^ provided a detailed structural representation with high confidence only for the conserved DNA polymerase core, covering 647 amino acids (aa) out of the full-length sequence of 1,306 aa for YerA41 DNAP I and 560 aa out of 1,175 aa for phiLo DNAP, including the palm, finger, thumb, and 3’-5’ exonuclease domains (**Suppl. Fig. 2d**).

### Domain organization in cryo-EM structures

We determined the cryo-EM structures of full-length DNAPs from the YerA41 and phiLo phages in multiple conformational states, including the apo form and DNA-bound complexes (**Fig. 2b-d, Fig. 3a-f, Suppl. Table 1, Suppl. Fig. 3-5**). In addition to the conserved 3’-5’ exonuclease core and characteristic palm, finger, and thumb domains of A-family DNAPs, there are several unique structural features specific to these enzymes. While some of these elements were identifiable as large insertions through sequence alignment (**Fig. 2a,c, Suppl. Fig. 1**), their function was revealed by the cryo-EM structures (**Fig. 2b,d**). Notably, both polymerases possess distinct insertions within the 3’-5’ exonuclease and thumb domains, designated as Insert 1 and Insert 2, respectively (**Fig. 2a,c**). In *YerA41* DNAP, Insert 1 spans residues 341 to 452 and consists of a four-helix bundle, augmented by two additional helices (**Fig. 2d**). Insert 2 spans residues 568 to 748 and is also composed of a four-helix bundle (**Fig. 2d**). In contrast, Insert 1 in *phiLo* DNAP is shorter, spanning residues 377 to 458, and contains three helices (**Fig. 2b**). Similarly, Insert 2 in phiLo is reduced in size but still consists of four helices arranged in a compact structure **(Fig. 2b).** These structural insertions create two extended arms located about 30 Å apart (in apo *YerA41* the distance between N610 and D390 is 33 Å and in apo *phiLo* between E574 and N410, this distance is 32 Å), flanking the DNA-binding site **(Fig. 2b,d).** Despite low conservation of these regions **(Suppl. Fig. 1)**, the structural positioning remains highly conserved between the both DNAPs structures **(Fig. 2b,d)**. A third structural element located at the C-terminal end of these phage DNAPs adopts a structure consisting of two β-strands and one α-helix, followed by a β-long-loop-β motif (**Fig. 2b,d – Suppl. Fig. 1**). This β-sheet is aligned with the β-sheet of the palm domain, effectively enlarging it and potentially contributing to its structural stability. The β-long loop-β structure forms a positively charged loop that is sandwiched between the thumb and palm domains which projects into the active site of DNAPs (residues 1272-1280 for YerA41 DNA polymerase and 1140-1156 for phiLo DNA polymerase) near the catalytic residues of the palm domain (**Fig. 4.b,d**). Despite the low sequence identity between the two polymerases, their structural similarity remains high, as indicated by a TM-score^39^ of approximately 0.71 if normalized by the length of phiLo DNAP after superimposition (**Suppl. Fig. 6c**).

**Figure 3.**
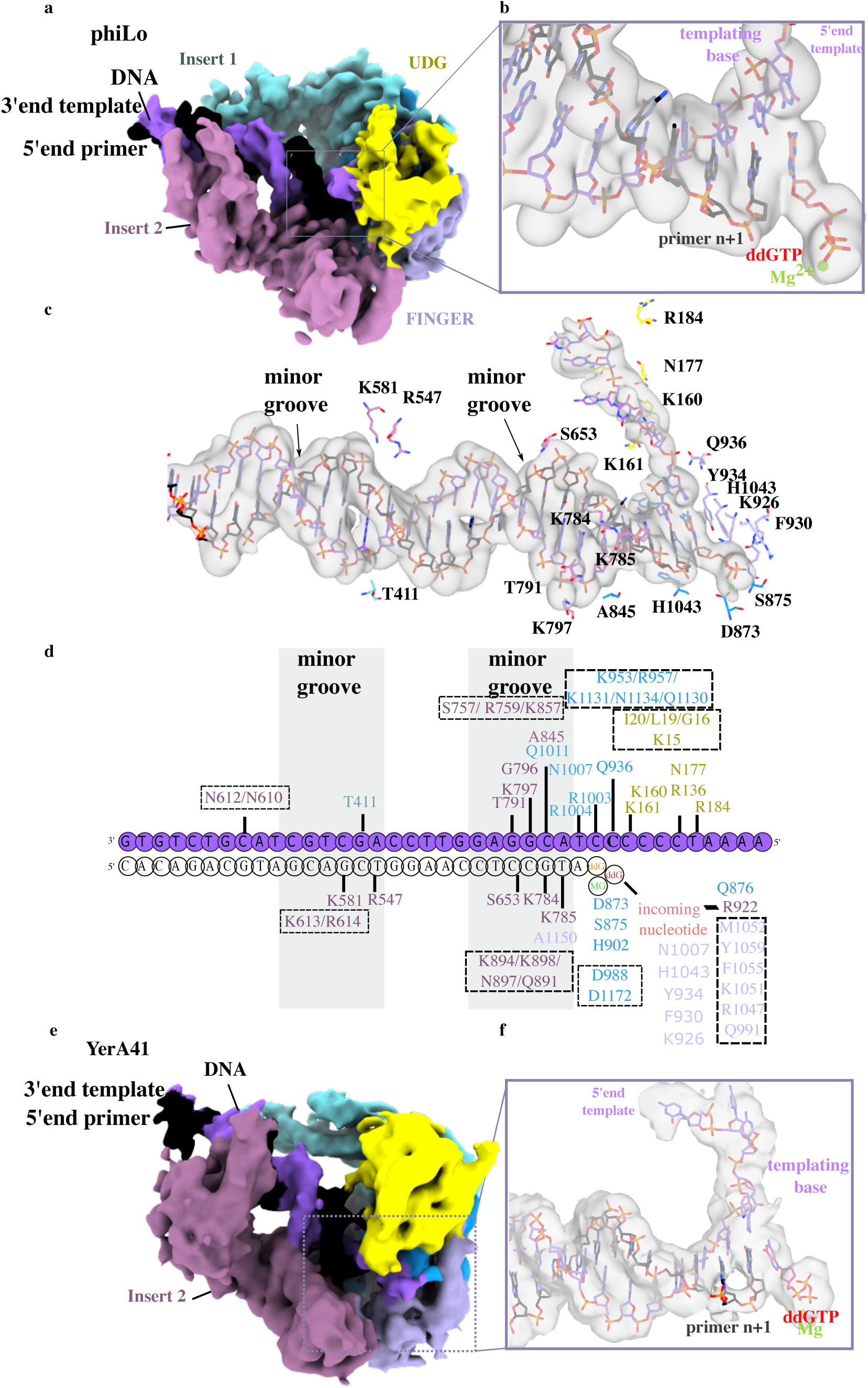
Cryo-EM structures of holoenzymes complexes. **a.** Cryo-EM map of phiLo DNAP as a complex with a DNA duplex substrate, that is shown in (c). The map is color-coded as in **Fig. 2**, with the primer strand in black and the template strand in purple. Insert 1 and Insert 2 enclose the DNA duplex on the 5’ side of the primer strand. **b.** Detailed view of the active site, illustrating base pairing and stacking interactions between bases in the electronic density represented in a transparent light grey map. The incoming nucleotide and Mg^2+^ ion are labeled as well as the 5’end template and the templating base. **c.** Electron density map in light grey for the DNA duplex, highlighting interactions between phiLo DNAP key residues, labeled in black, and the minor groove of the DNA. **d.** DNA duplex used in this cryo-EM structure. The template DNA substrate is color-coded in purple. Two deoxycytosine bases serve as template bases, complementary to the incoming ddGTP nucleotide. Key residues from phiLo DNAP involved in the DNA duplex stabilization are indicated with a domain-wise color coding, while key residues from YerA41 DNAP are squared. **e.** Global view of the cryo-EM map (color-coded by domain) of YerA4 DNAP in the ternary complex conformation. **f.** Electron density in transparency surface mode and stick representation of the DNA duplex substrate and its 5’ template DNA strand extension, also showing the incoming nucleotide and a magnesium ion (green).

**Figure 4.**
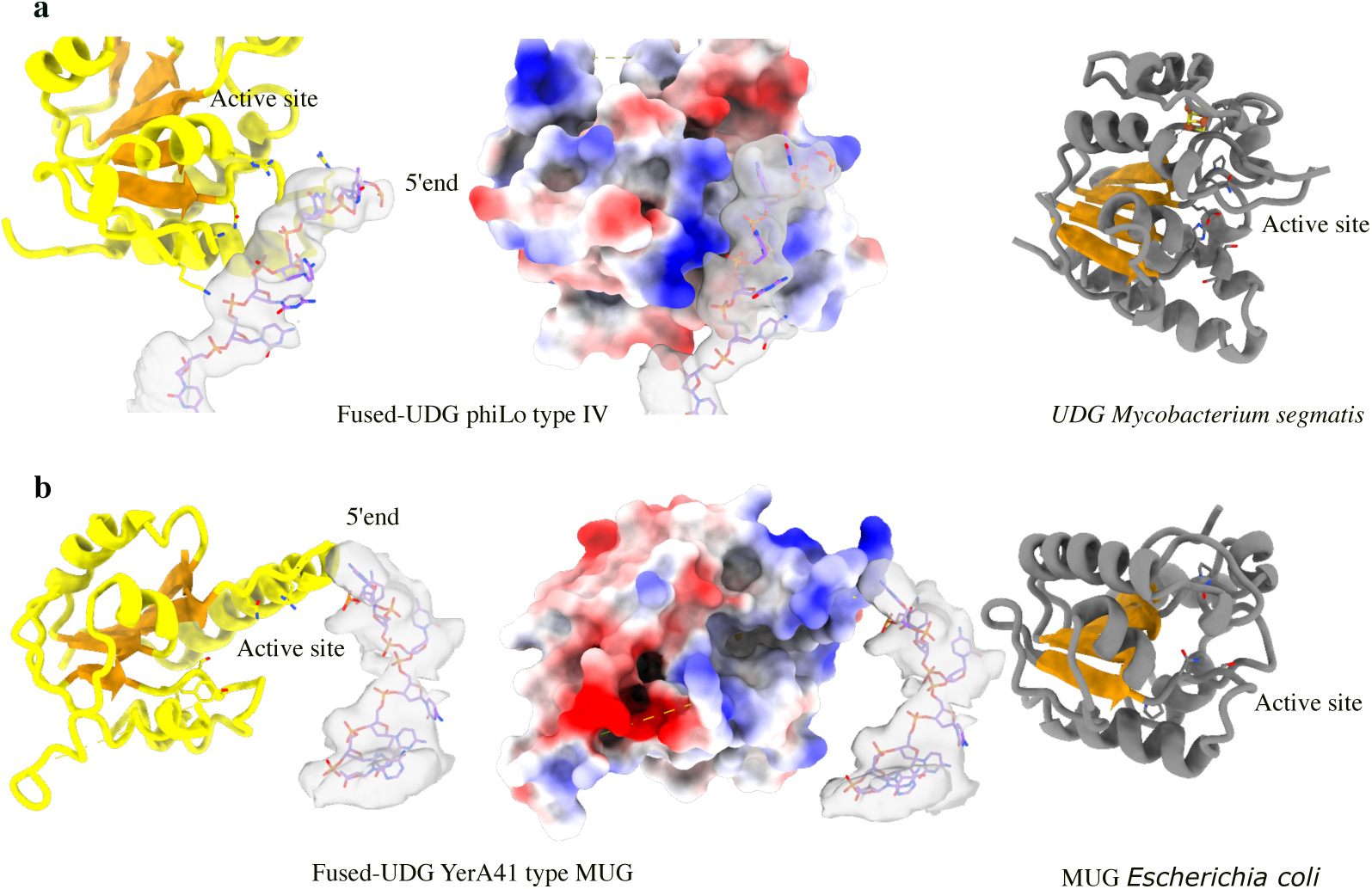
Template strand 5’-end stabilization by the positively charged UDG domain fused to SP15-like DNAPs. **a. Left panel:** The type IV UDG domain (yellow) from phiLo DNAP interacts with the 5′ end of the template strand, shown in backbone representation with its experimental electron density overlaid as a transparent surface. The catalytic residues from motifs A and B, identified in the MSA (**Supp. Fig. 1**), are shown in stick representation. **Middle panel:** Electrostatic potential projected on the surface (positive region are colored in blue, negative region are colored in red) of the phiLo UDG domain, highlighting the 5′ end of the template strand (also shown as a transparent surface). **Right panel:** The structure of *Mycobacterium smegmatis* (Msm) UdgX (gray) is displayed in the same orientation, with the core β-sheet in orange. Specific residues involved in binding ssDNA and removing the dU base (in stick representation) are highlighted (PDB ID: 6IOD). **b. Left panel:** The type II UDG domain (yellow) from YerA41 DNAP is positioned similarly on top of the finger domain but is slightly shifted toward the 3′-5′ exonuclease domain. The 5′ end of the template is drawn in a bones representation, together with its experimental electron density in surface mode. Catalytic residues are displayed in stick. **Middle panel:** Electrostatic potential projected on the surface (positive and negative regions are colored in blue and red, respectively) of the YerA41 UDG domain, highlighting the 5′ end of the template strand (also shown with its transparent surface). **Right panel:** X-ray structure (in grey) of the *E. coli* MUG enzyme in the same orientation (PDB ID: 4MWI). The core internal β-sheet is in orange.

These three insertions serve as defining signatures of SP15-like polymerases, marking them as a novel subfamily within A-family DNA polymerases. While they do not form a distinct cluster in the CLANS 3D map, they appear as a radial extension away from the central cluster in **Fig. 1a**. The addition of similar sequences in future analyses is likely to reinforce this trend.

### Conformation of SP15-like DNAP poised for DNA primer elongation

We then determined the cryo-EM structures of both phiLo and YerA41 DNAP in the presence of primer-template DNA substrates and dideoxyguanosine (ddGTP) as the incoming nucleotide (as shown in **Fig. 3**) in order to trap the ternary complex. These structures, solved at 3.7 Å and 4.5 Å resolution, respectively, captured the enzymes in the elongation state (**Fig. 3a-e**). The observed electron density allowed us to confidently model the primer-template duplex and revealed the precise positioning of the incoming nucleotide, ddGTP, at the 3’-OH end of the primer (**Fig. 3b, 3f**). Structural analysis revealed a single incorporation event of ddGTP, resulting in a +1 extended primer terminus that lacks a 3’- hydroxyl group. Additionally, we observed a second ddGTP molecule occupying the active site, that forms a Watson-Crick base pair with the templating base but has yet to be incorporated into the primer strand. This interaction is stabilized by specific residues in the O-helix of the finger domain (**Fig. 3c,d and Suppl. Fig. 7b-c**) with the polymerase adopting a closed conformation of the active site. This structural change is evident from the 14.5 Å shift of the O-helix compared to the apo form (measured between N921 residues in the apo and holo conformations of phiLo). Additionally, a Mg^++^ metal ion is observed in the electronic density, coordinated by the ⍺ and β phosphates of ddGTP and the residue D873 in phiLo (**Fig. 3c,d**), and D988 in YerA41 (**Fig. 3d**), which is typically the position of metal “B” in the two-metal ion catalysis mechanism described in canonical DNA polymerases^40,41^. Furthermore, the ddGTP sugar moiety packs against F930 (phiLo) and F1055 (YerA41) from the O-helix, which is critical to discriminate deoxynucleotides against ribonucleotides by providing a steric clash with the ribonucleotide-specific 2’-OH group^42^.

### Insert 1 and Insert 2 are involved in multiple interactions with DNA in holoenzyme complexe

Beyond the well-characterized roles of the finger and palm domains in DNA binding and structural rearrangement during elongation (**Fig. 3b,g**), we could identify additional stabilization mechanisms mediated by Insert 1 and Insert 2, which enclose the double-stranded DNA (dsDNA) following the replication site. These structural elements undergo subtle yet functionally significant conformational adjustments to reinforce the interaction with the primer-template duplex (**Fig. 3c,d; Suppl. Fig. 7c**). Residues within Insert 2 establish direct interactions with the dsDNA backbone, primarily through contacts in the minor groove of the DNA, while the major groove remains solvent-exposed (**Fig. 3e**). In phiLo DNA polymerase, Insert 2-mediated contacts involve N649, P650, S651, S652, K581, and Q585 (interestingly all these residues are well conserved in the SP15-like family members, see **Suppl. Fig. 1**), while Insert 1 contributes additional stabilizing interactions notably via T411, which stands out as a well conserved position in Insert1 (**Suppl. Fig. 1**) even it is fully exposed residue. These findings highlight a complementary role for Insert 1 in dsDNA stabilization, in conjunction with Insert 2. Notably, Insert 2 undergoes a transition from a position ∼5 Å away to an enclosed conformation, wrapping around the DNA upstream of the active site (**Fig. 3b**). In YerA41 DNA polymerase, the OP1 and OP2 groups of the dG20 base form hydrogen bonds with N610 and N612 in Insert 2, whereas dG9 and dT10 interact with R759 in the thumb domain via hydrogen bonding (**Suppl. Fig. 7b,c**). Additional stabilizing interactions involve G611, F696, F755, S757, and T758. Interestingly, no direct Insert 1 contacts were detected, although it is very close to the DNA path. We hypothesize that Insert 1 may undergo dynamic rearrangements during elongation, transiently engaging with the dsDNA backbone to further stabilize the polymerase-DNA interaction.

These distinctive insertions appear to enhance the stability of the ternary complex, potentially increasing polymerase processivity—a characteristic feature of SP15-like polymerases.

Multiple sequence alignments (MSA) across this family of polymerases indicate that, while the precise length and sequence composition of these inserts differ, their spatial localization around the DNA remains conserved (**Suppl. Fig. 2a**). In contrast, SPO1-like polymerases exhibit only a truncated region at the position of Insert 2, which folds into two helices integrated within the thumb domain (**Suppl. Fig. 2b**). The absence of Insert 1 and of a fully developed Insert 2 in these polymerases likely contributes to their distinct structural clustering away from SP15-like polymerases. Additionally, no insertions were observed within the 3’-5’ exonuclease domain, reinforcing the hypothesis that Insert 1 and Insert 2 have evolved as polymerase-specific DNA stabilization clamps.

Interestingly, the spatial positioning of Insert 2 relative to the dsDNA backbone is reminiscent of the thioredoxin binding mode assisting in T7 replication^43^. This suggests that Insert 2 may serve a role analogous to thioredoxin-mediated polymerase processivity enhancement, with increased replication efficiency. Expanding beyond bacteriophages, similar stabilizing cofactors are observed in DNA polymerases from viruses infecting humans, including Monkeypox virus, Herpesvirus, and African swine fever virus. Notably, viral replication factors such as H5-H5’ and A22 are thought to function analogously to thioredoxin^44^.

### A guiding channel for the 5’ end of the template strand toward the UDG domain active site

In the SP15-like A-family DNAP structures, the N-terminal UDG domain is positioned between the finger and exonuclease domains (**Fig. 2**). Due to missing electron density in certain regions, only specific segments could be traced: residues 1–91 and 141–190 in YerA41 DNAP and residues 39–90 and 119–210 in phiLo DNAP. Despite this incomplete assignment and model building in the electron density, sequence analysis clearly classified the UDG domain in YerA41 as type II and in phiLo as type IV. In phiLo DNAP, the UDG domain adopts a somewhat distinct spatial arrangement, positioned closer to the thumb domain, whereas in YerA41, it is located near the 3’–5’ exonuclease domain. The UDG domain in phiLo interacts with nucleotides C-2 to C-4 that are clearly visible in the electron density, while nucleotides spanning positions C-5 to A-9 remain unresolved (**Fig. 4a left panel**). Analysis of the electrostatic potential of the UDG domain reveals a large positively charged surface patch, compatible with the binding site of the ssDNA (**Fig. 4a middle panel**). Structural comparison with *Mycobacterium smegmatis* UDG (MsmUDG) revealed a conserved fold, with the characteristic LVGENP and HSS sequence motifs preserved as LVGEAP and HSF in phiLo DNAP, suggesting functional conservation (**Fig. 4a right panel**). While the stabilization of the ssDNA 5’end template is clearly seen, the template doesn’t reach the active site of the UDG domain as observed into the crystallographic structure of the MsmUDG homologue (**Fig. 4a**). Of note, the template strand does not contain a dU base in this experiment. In YerA41 DNAP, the UDG domain exhibits the canonical fold of the UDG family, characterized by a central four-stranded β-sheet flanked by helices (**Fig. 4b**). This structure closely resembles *Escherichia coli* uracil DNA-glycosylase (MUG), where the conserved GINPG and NPSGLS sequence motifs form the active site responsible for uracil excision and abasic site generation, preserved as GYVN and MYSLM in YerA41 DNAP (**Fig. 4b right panel**). In the DNA template strand, nucleotides from −3 to 25 were modeled in YerA41 DNAP ternary complex, revealing a channel formed by K15, G16, L19, and I20, which interacts with C-3 at the 5’ end of the template (**Fig. 4b**). Although the rest of the 5’ end of the ssDNA could not be modeled within the UDG pocket, likely due an intrinsic high mobility, sequence analysis identified the conserved GYVN and MYSLM motifs in corresponding secondary structures (**Fig. 4b, Suppl. Fig. 1**). As observed in phiLo DNAP, the 5’end of the ssDNA is not bound within the active site of the UDG domain, likely due to the absence of an uracil base at the 5’end of the template. To further investigate this, a DNA substrate containing uracil was used in the following part of the study.

We note that the positioning of the UDG domain in these polymerases is similar to its location in the trans-complex of *Escherichia coli* DNA polymerase I with the endonuclease FEN1^45^, which ensures coordinated nuclease and strand displacement activities (**Suppl. Fig. 8**). Strikingly, it has been proposed that FEN1 and the UDG family share a common evolutionary ancestor^23^, and the similar localization of both domains in a DNAP context supports this hypothesis ^23^.

### Mechanism of dU excision prior to replication

The specific activity of base excision when encountering deoxyuracil (dU) bases was tested on SP15- like DNAPs using a DNA duplex containing dU (**Fig. 5a**), followed by treatment with the endonuclease APE1. The appearance of a 14-nucleotide fragment, as detected by Cy5 fluorescence, indicates that phage DNA polymerases with UDG-like domains do excise dU, with APE1 subsequently cleaving the resulting abasic site (**Fig. 5b).** In the presence of dNTPs, we observed primer extension with a 34- nucleotide fragment, demonstrating lesion bypass activity (**Fig. 5c**), that will be described in details in the following section.

**Figure 5.**
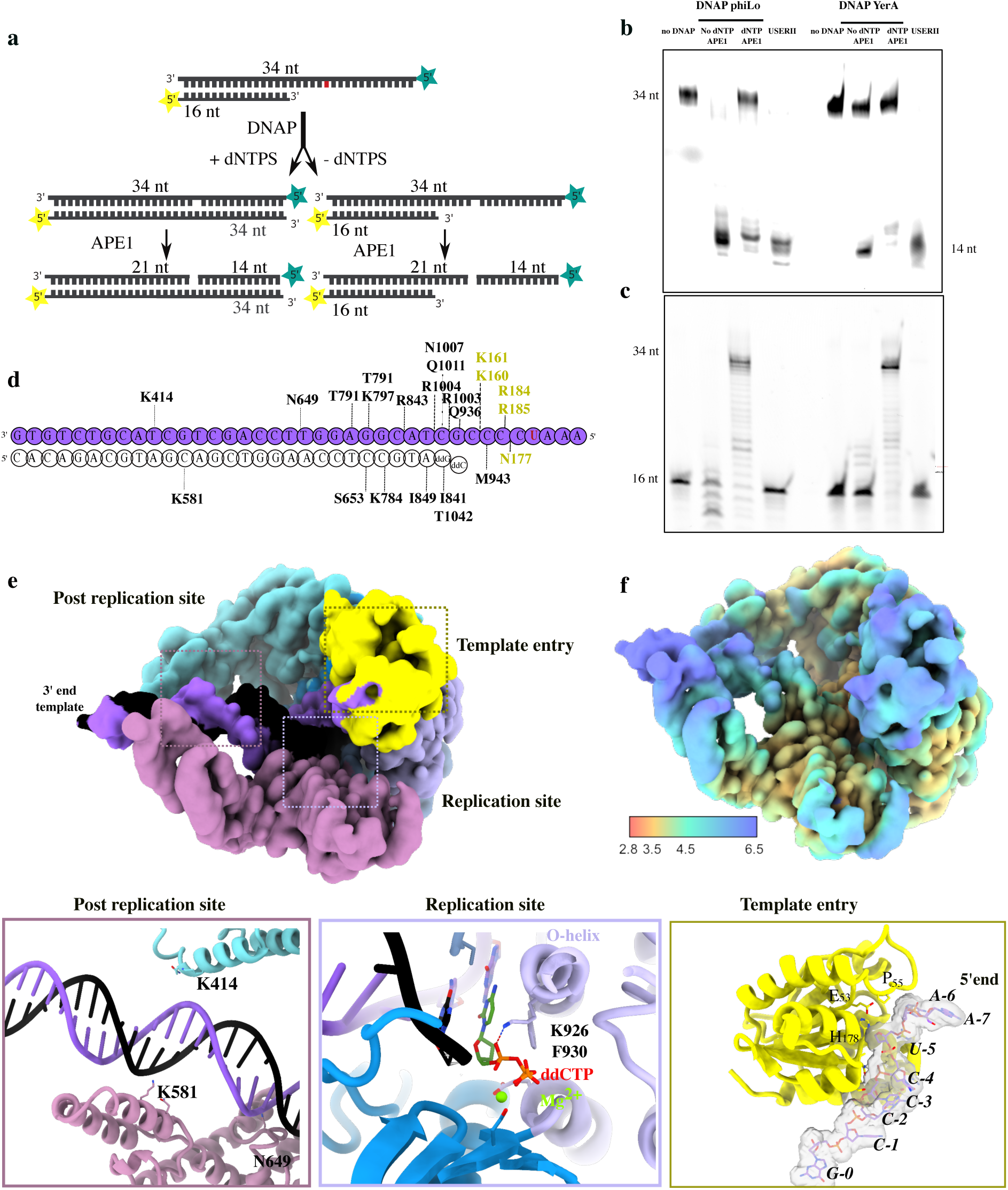
Structural basis for the excision of dU in the 5’ end of the DNA template strand by DNAPs. **a.** The primer extension activity of DNAP YerA41 and DNAP phiLo was evaluated using a 5’-FAM-labeled primer DNA substrate and a 5’-Cy5-labeled dU-containing template DNA strand. The FAM label is in yellow and the Cy5 label is represented as a blue star. The dU base is in red. The lengths of the corresponding fragments are indicated. This assay was followed by APE I endonuclease activity, which removes the AP site generated by the UDG domain of the DNAPs. **b.** Denaturing gel (8 M urea, 16% acrylamide) scanned for the Cy5 fluorescence indicating the length of the 5′- labeled template strand after elongation by DNAPs. A control with USERII is provided. **c.** Denaturing gel scanned for FAM fluorescence, probing the progression of DNA synthesis (length of the primer strand) in different conditions, as indicated on top of the lanes. **d.** The DNA duplex used in this cryo-EM structure. The template DNA substrate is color-coded in purple. The primer strand is in black. The first two templating bases are a deoxycytosine and a deoxyguanosine base, complementary to the incoming ddGTP and ddCTP nucleotides added in solution. Key residues involved in stabilizing the DNA are indicated. **e.** The cryo-EM density map of phiLo DNAP bound to the DNA duplex substrate, with domain colors corresponding to **Fig. 1a**. Three zoomed-in views highlight the active site of phiLo DNAP interacting with the DNA substrate and the incoming ddCTP nucleotide (bottom middle, Replication site) and the interactions of Inserts 1 and 2 with the DNA duplex (bottom left, Post-replication site) and the UDG domain (bottom right, Template entry). The latter view illustrates key amino acids residues (numbered and depicted as sticks) stabilizing the DNA template strand near the 5′ end, whose numbering is shown in black together with its electron density in a grey surface. The −5 position that should contain the dU base is flipped and localized just at the entry of the active site near catalytic residues H178, P55 and E53.The absence of density for the dU base itself is indicative of the activity of the UDG domain, which generates an AP site (**b**). **f.** The cryo-EM map colored according to the local resolution scale bar on the bottom.

We then solved by cryo-EM a third structure of phiLo A-family DNA polymerase in the presence of a primer-template containing dU at the 5’ end (**Fig. 5d-f**). A high resolution map reaching locally 2.1 Å was obtained, allowing for precise modeling of the DNA substrate. As in the previous structures, ddCTP and ddGTP that are complementary to the first two templating bases were added to capture the elongation mode. The O-helix indeed adopts a closed conformation stabilizing the second incoming nucleotide, ddCTP (**Fig. 5d,e**). Residues K926 and F930 and a coordinating Mg²⁺ interact with the triphosphate moiety, while the base of the incoming ddCTP makes Watson-Crick interaction with the templating base (**Fig. 5e**). As expected, we observed density corresponding to the 5’ end of the DNA template strand near the active site of the UDG-like domain, and the dU base is positioned five residues downstream of the DNA polymerase replication site (**Fig. 5e**). Key residues, including K161, R185, N177, and F181, primarily stabilize the DNA backbone between nucleotide positions C-1 and U-5. The electron density map provides clear evidence of base stacking from C-1 to C-4; however, the absence of density for the dU base itself suggests that the UDG-like domain has already been excised from the DNA template. This observation demonstrates that the SP15-like members DNAP have two distinct enzyme activities and underscores the role of the UDG-like domain in excising dU.

Interestingly, in the replicative complex of the Mpox virus, the B-family DNA polymerase physically interacts with the E4 protein, encoded in trans in the Monkeypox virus genome and belonging to the UDG family^28,29,44,46^. The UDG protein also scans the 5’ end of the template ahead of the DNA polymerase’s replicative pocket, highlighting a direct connection between dU repair and DNA replication.

### Role of the C-terminal domain

To gain further insight in the mechanism of dU excision ahead of the replication site, we tested the ability of SP15-like DNA polymerases to bypass an abasic site. In the presence of dNTPs, phage DNA polymerases could replicate a single-stranded DNA template containing an abasic site analogue, tetrahydrofuran (THF), immediately followed by a T base, confirming their lesion bypass activity (**Fig. 6a**). Both DNA polymerases from YerA41 and phiLo were able to incorporate a nucleotide opposite the THF site. In terms of selectivity, these polymerases preferentially incorporated dATP, which is consistent with the ‘A-rule’ commonly observed in other DNA polymerases^47,48^ (**Fig. 6b**). For comparison, the lesion bypass mechanism of Taq DNA polymerase (PDB ID 3LWL) involves residue R587 from the thumb domain and Y671 from the O-helix. Y671stabilizes ddATP in front of an AP site and participates to the ‘A-rule’ mechanism in bypassing abasic sites, while R587 stabilizes the incoming adenine mediated by a water molecule (**Fig. 6c)**. In the cryo-EM structure, these residues of the thumb domain were not positioned near the replication site, because these DNA polymerases undergo a major structural rearrangement of the thumb domain compared to conventional DNA polymerases. Instead, the β-long-loop-β structural motif from the C-terminal domain that protrudes into the active site may play the role of R587 in Taq polymerase by providing positively charged residues (K1273 - H1274 from *YerA41* and K1148 – K1149 from *phiLo*) (**Fig. 6d,e**). The CTD is conserved among all SP15-like DNA polymerases as well as the β-long-loop-β loop structural motif providing positively charged residues (**Suppl. Fig. 1**).

**Figure 6.**
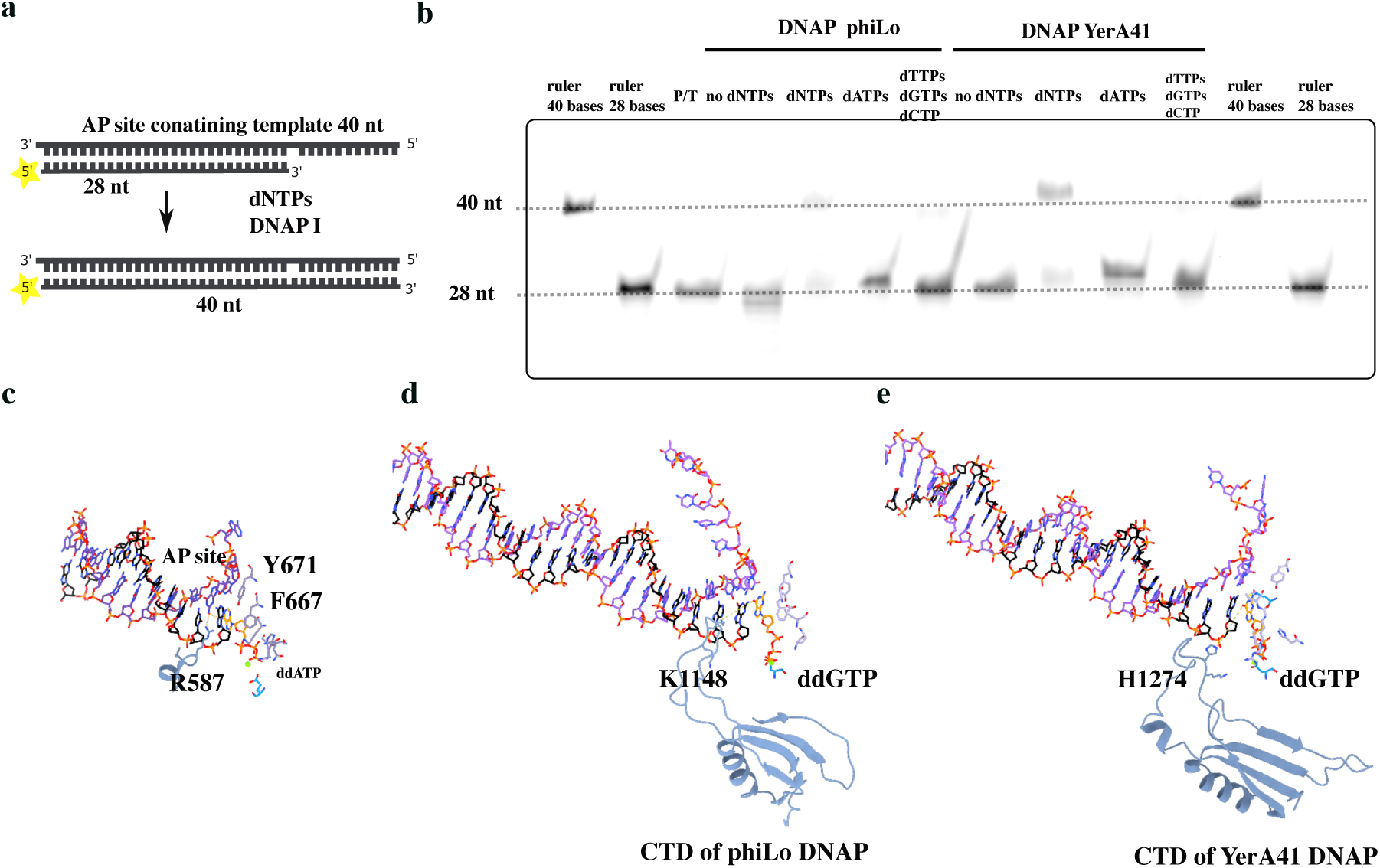
DNA replication through abasic sites: a possible structural role for the C-terminal domain. **a.** Bypass of an abasic site by DNAP 144 from YerA41 and DNAP 158 from phiLo assessed using a 5’-FAM- labeled primed substrate (yellow star) base-paired to a DNA-template strand containing a THF (AP) site in the templating position. The lengths of the strands are indicated. The THF site is indicated by the absence of a base. **b.** Primer extension test across a THF site visualized by PAGE with FAM fluorescence under different conditions: no dNTP, a complete dNTP mix, dATP alone, and a combination of dGTP, dCTP, dTTP. **c.** Close-up view of the DNA substrate in KlenTaq DNA polymerase ternary complex structure (PDB ID 3LWL) with a template DNA strand (in purple) containing an abasic (AP) site and the incoming nucleotide. The primer strand is in black. The incoming nucleotide (ddATP, in stick representation) is stabilized by Y671 and F667 from the O-helix along with R587 (in blue). Y671 mimics the absent templating nucleobase while R587 interacts with a water molecule to stabilize the ddATP incoming nucleotide. The catalytic aspartate is in cyan and the magnesium is in green (also shown in panels **d** and **e**). **d.** Structure of phiLo DNAP, showing only the DNA duplex and the CTD extension represented in a cartoon mode. A closed-up view of the β-long loop-β structure (in blue) in the C-terminal domain shows the KK motif that stabilizes the incoming nucleotide. Residues stabilizing the ddGTP incoming nucleotide are also represented. **e.** Structure of YerA41DNAP, showing only the DNA duplex with the template strand in magenta and the primer strand in black. A close-up view of the β- long loop-β structure (in blue) in the C-terminal domain (CTD) highlights a KH motif (in sticks) that stabilizes the incoming nucleotide in the absence of the templating nucleobase, playing the same role as R587 in KlenTaq.

In DNA polymerase theta, two specific insertions into the palm domain were probed and described to be important for the bypass of the THF base, but they differ from the C-terminal domain described here^49^.

### Interplay between DNA synthesis and DNA repair in SP15-like phages

To address the fate of abasic sites generated by the UDG domain prior to replication, we analyzed the genes encoded in the phage genomes, focusing on those involved in AP site repair. We identified the genes encoding DNA polymerase X (gene 158 for YerA41) and DNA ligase (gene 127 for YerA41 and gene 188 for phiLo) (**Fig. 7a**). All members of the SP15-like DNA polymerase family contain genes encoding both DNA polymerase X and DNA ligase within their genomes, except for the phiLo genome where the DNA polX could not be detected (**Fig. 7a**). Interestingly, genomes corresponding to SPO1- like polymerases lack genes encoding DNA polymerase X, highlighting a unique characteristic of SP15-like phage genomes. There seems to be no specific apurinic endonucleases (APE) in the genomes studied but it has been reported that some viral DNA polymerase X enzymes do possess both APE and gap-filling activities^50–53^. To test the repair activity of these proteins, activity assays were performed using DNA substrates containing a deoxyuridine (dU), a tetrahydrofuran (THF) abasic site analog, and a single-nucleotide gap (**Fig. 7d**). In the presence of DNA polymerase I from *YerA41*, we observed the cleavage of a DNA strand containing a single deoxyuracil (dU), resulting in a 15-nucleotide fragment upon the subsequent addition of DNA polymerase X from *YerA41* **(Fig. 7c Left panel)**. In contrast, no cleavage product was detected in the absence of A-family DNA polymerase, suggesting the absence of an abasic site formation. As a control, we employed a commercial UDG along with two distinct AP endonucleases: endonuclease IV (endo IV) and APE1. APE1 generates a 3’-unsaturated aldehyde intermediate fragment, while endo IV produces a 3’-OH fragment. Notably, the DNA polymerase X generated a 3’-OH product, consistent with the activity profile of endo IV **(Fig. 7c Left panel)**. To further validate the AP endonuclease activity of DNA polymerase X, we conducted an AP endonuclease activity assay using a DNA duplex containing a THF site, as described in **Fig. 7d**. We confirmed that DNA polymerase X cleaved the substrate to yield a 15-nucleotide fragment **(Fig. 7c Middle panel)**. Finally, we investigated the gap-filling activity in conjunction with DNA ligase I from *YerA41*. We assembled a DNA duplex containing a gap (**Fig. 7d**) and performed a ligation assay in the presence of DNA polymerase X and DNA ligase I. This experiment demonstrated that DNA polymerase X and DNA ligase I successfully generated a 34-nucleotide fragment, indicative of complete DNA repair and the formation of a nick-free DNA product **(Fig. 7c Right panel)**.

**Figure 7.**
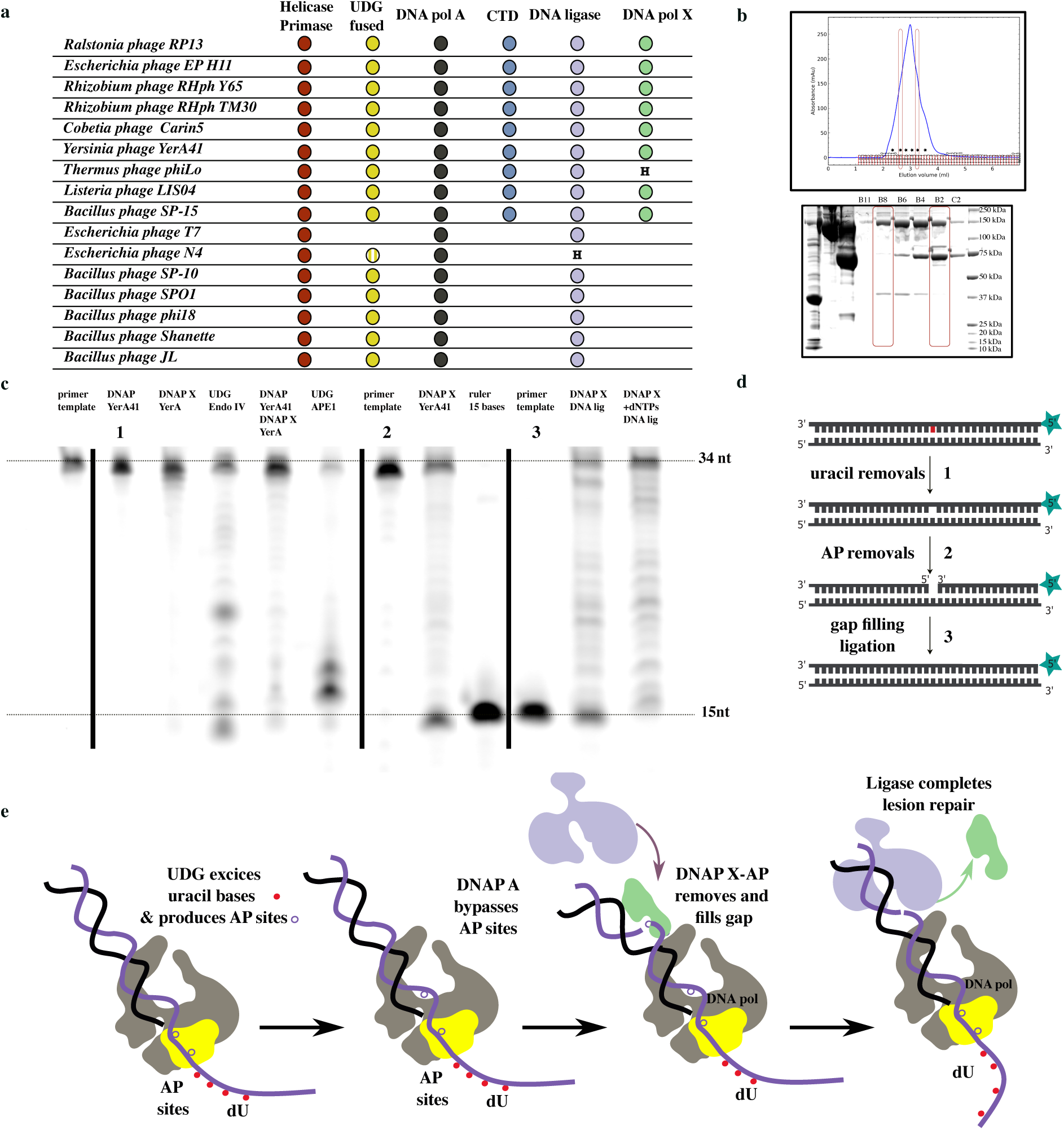
Coupling of DNA replication and DNA repair mechanisms in presence of dU bases in the phage DNA. **a.** Co-occurrence of DNA polymerase I (gray circle) with the UDG domain (yellow circle), CTD domain (blue circle), DNA ligase (purple circle), DNA polymerase X (green circle), and helicase/primase (red circle) in SP15- like members is compared to their presence in T7, N4, and SPO1-like phages. In contrast, SPO1-like members, which lack the CTD (see AlphaFold models of DNA polymerases in **Suppl. Fig. 2**), possess a DNA ligase but do not have a recognizable DNA polymerase X. N4-like phages contain only a A-family DNA polymerase with a UDG domain that lacks catalytic residues and the CTD. The DNA ligase is present in the host genome (H label). For phiLo, the gene encoding DNAP X is not found in the phage but there is one in the host *T.thermophilus* HB8 (H label). **b.** Gel filtration (top panel) of YerA41 DNAP in the presence of X-family DNAP, DNA ligase, and the DNA substrate oSM0072/oSM0074. The elution fractions were analyzed by SDS-PAGE (bottom panel). The elution profile revealed two distinct peaks: one corresponding to the co-elution of YerA41 DNAP and X-family DNAP, while DNA ligase eluted separately at a later stage, enclosed by a red rectangle. **c.** Left: Reconstitution of step 1 in (d). Activity assay of A-family DNAP coupled to X-family DNAP from YerA41 on a primer-template containing dU base(s). A cleaved product is observed in the presence of both A- family DNAP and X-family DNAP from YerA41. Control experiments were conducted with commercially available UDG proteins EndoIV and APE1 from NEB. DNA polymerase X exhibits AP endonuclease activity similar to that observed for EndoIV, leaving a one-nucleotide gap with a 3’ hydroxyl and 5’ deoxyribose phosphate (dRP) termini, while APE1 cleaves the abasic site, producing a 3’ α-β unsaturated aldehyde and a 5’ phosphate termini. Middle: Reconstitution of the step 2 in (d). Activity assay of X-family DNA polymerase from YerA1 in front of a THF site (mimicking an AP site) containing DNA. A cleavage product is observed, indicating intrinsic AP endonuclease activity in the polX protein. Right: Reconstitution of the step 3 in (d): Activity assay of the X- family DNAP coupled to the DNA ligase from YerA4 to test the gap-sealing function of these enzymes. In the presence of X-family DNAP and dNTPs, the DNA ligase from YerA41 generates a DNA product with a length of 34 nucleotides. Without dNTPs, the DNA ligase is able to seal the extremities of DNA across the gap. **d.** Different steps in the processing of a dU-containing DNA substrate used for the activity assays described in (**c**), involving the phage DNA polymerase fused to an active UDG domain, a phage DNA polymerase X and a phage DNA ligase. An uracil base (red) is positioned at +16 nt on the template, as well as a THF site. The final substrate is a DNA duplex encompassing a one-nucleotide gap with a substrate of 15 bases. **e.** Steps of DNA replication coupled to the DNA repair mechanism generalized for SP15-like members. Four different steps can be identified. The X-family DNAP is in green, and the DNA ligase is in light purple.

To explore the possible formation of a supramolecular repair complex, we performed size exclusion chromatography using reconstituted complexes of A-family DNAP, PolX, and DNA ligase. Two main peaks were observed with a retention volume of 2.9 mL and 3 mL, respectively. SDS-PAGE analysis of the different elution fractions revealed that the DNAP A and DNAP X proteins co-elute within the same peak, indicating the formation of a complex in the presence of a substrate DNA duplex ( **Fig. 7b**). This suggests that A-family DNAP and DNAP-X work together in a coordinated manner. The second peak corresponds to the DNA ligase, suggesting that the recruiting of this factor comes later.

In line with all these results, we propose a model (**Fig. 7e**) in which the UDG domain scans the 5’ end of the template ahead of the replication site. When a dU base or modified base is encountered, the active UDG domain excises the lesion, leaving behind an abasic (AP) site. The DNA polymerase’s active site can subsequently bypass the AP site lesion, enabling continuation of replication. Following replication, DNA polymerase X, which exhibits AP endonuclease activity, processes and fills in the resulting gap. Finally, the DNA ligase completes the repair process by sealing the nick in the DNA. Altogether, our data suggest that the replication machinery and the AP site repair pathway may work in a streamlined process, effectively bypassing the host’s defense mechanisms that would otherwise excise dU-containing phage DNA. The co-occurrence of these genes strongly supports the hypothesis that these phages have evolved an integrated mechanism to coordinate DNA replication and repair (**Fig. 7a-e**).

## Discussion

This study reveals a mechanism by which certain phages with highly modified DNA integrate both DNA elongation and dU-lesion recognition and excision within a single enzyme. We show that these large A-Family phage DNAPs, when fused with a UDG domain, are capable of excising uracil, generating abasic (AP) sites, and bypassing these lesions while copying phage DNA (**Fig. 7**). This suggest that the *timing* of base excision repair (BER) in theses phages - and possibly some DNA viruses - differs from both eukaryotic and bacterial repair systems. The structural analysis shows how the UDG domain interacts with the DNA template ahead of the replication site, excising uracil residues while the polymerase domain simultaneously synthesizes the nascent DNA strand. By removing dU sites during replication, this system acts as counter-measure against bacterial defense systems, such as increasing the concentration of dUTP in the cell to compete with the incorporation of 5-hmdUTP. One advantage to have the UDG domain *in cis*, rather than *in trans* would to avoid inactivation by a small protein (about 80 aa) called uracil-DNA glycosylase inhibitor protein (Ugi) that is present in some microorganisms by concealing its interaction surface^23^. Ugi specifically binds and inactivates UDG enzymes, thereby blocking the base excision repair (BER) pathway^21,54^.

At least two SP15-like phage genomes contain hyper-modified thymidines in their genomes, SP-15 and YerA41. This could make the job of the UDG easier, taking advantage of the large difference between the uracil and the hyper-modified thymine compared to the small difference between uracil and thymine as position 5 of the pyrimidine ring. In addition, the analysis of contact points between the SP15-like DNAPs and the DNA duplex show that they mostly involve the phosphate backbone in the minor groove of the DNA duplex, leaving the major groove completely accessible to the solvent. This structural arrangement tentatively explains the accommodation of highly glycosylated DNA within the polymerase active site. However, experimental studies using glycosylated YerA41 DNA as a substrate are needed to fully establish the molecular adaptations of SP15-like DNA polymerases to highly modified DNA. The translesion synthesis activity (TLS) of some A-Family DNA polymerases has long been studied, especially in DNA polymerase theta^49^. The present structural studies suggest that the CTD domain of SP15-like DNAPs play a role in TLS across abasic sites. Specifically, the β-long-loop-β structure, which introduces several positively charged residues near the 3’-OH primer end and the incoming nucleotide base, may function similarly to the arginine (R587) residue in Taq polymerase to facilitate the bypass of abasic sites (**Fig. 6c-e**) on the template^55^. Additionally, previous studies have reported that the A-Family DNA polymerase YerA41 can initiate replication in a primer-independent manner^8^. It is possible that this function may be shared by other members of the SP15-like cluster and that the β-long-loop-β structure itself could contribute to primer-independent replication perhaps through the sequential insertion-retraction of this long loop. The CTD domain was not well predicted in DNAP YerA41 by AlphaFold2 (**Suppl. Fig. 2d**), whereas it was well predicted in DNAP phiLo. Overall, this domain appears challenging to model using prediction tools based on sequence analysis, suggesting that it may also be present but still undetected in other DNA polymerases.

Structural analysis reveals two large insertions that interact with double-stranded DNA (dsDNA), providing insights into a mechanism that enhances DNA replication processivity. Rather than forming a ring-like structure, these insertions stabilize DNA through their positively charged surfaces, resembling the thioredoxin domain found in the replicative T7 DNA polymerase^43^. In T7 DNA polymerase, this site is known to interact with the primase-helicase complex at the replication fork^56–58^ (**Fig. 8a,b**), suggesting a similar interaction between the replicative helicase encoded in SP15-like phages and their DNA polymerase. Based on this hypothesis, we can propose a model of the replisome in these phages, comparing it to the well-characterized replication machinery of T7 phage^56^. We take it that SP15-like phages utilize a single replicative DNA polymerase for both the leading and lagging strands, as only one replicative DNAP is encoded in their genomes.

**Figure 8.**
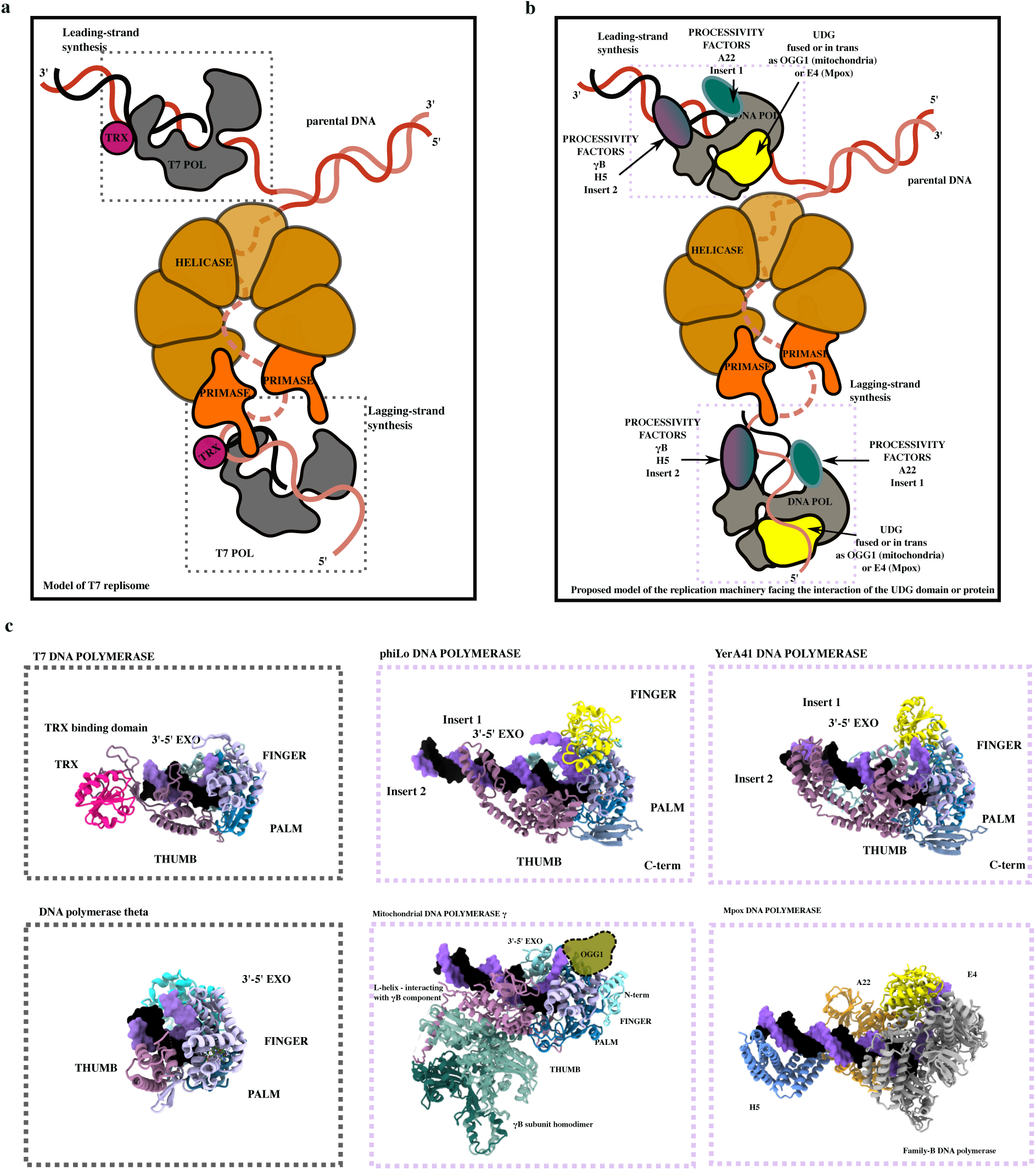
Comparison of A-family DNA Polymerase from YerA41 and phiLo Phages with T7 DNAP and its replisome, Human Mitochondrial DNAP γ and Human Viral Replicative DNAP. **a.** Schematic view of the the cryo-EM T7 replisome^75^, consisting in a complex of T7 DNA polymerase (in grey, with the thioredoxin TRX in magenta) with DNA (red) and the hexameric helicase (brown) at the lagging strand synthesis side (PDB ID 6N9V) as a complex with the primase (orange). **b.** Proposed model of of a T7-like replisome using SP15-like DNA polymerases, with a UDG protein (yellow) interacting with the DNA polymerases. This should be valid also for the Mpox DNA polymerase holoenzyme and its UDG domain in trans. In mitochondria, OGG1, a UDG protein known to work together with mtDNA polymerase *γ*, can be placed also in this context, although the mtSSB protein would be also necessary to fully understand the mitochondrial replisome. **c.** Zoomed-in panel showing the T7 DNA polymerase structure in complex with the thioredoxin protein (PDB ID 1ZYQ), the human Pol theta DNAP domain (PDB ID 8GD7), the DNA polymerases of YerA41 and phiLo, shown in the same orientation as the DNA polymerase *γ* (PDB ID 8V55) and DNA holoenzyme of Mpox, with a similar localization of the UDG domain and E4 of the Mpox virus (PDB ID 8WPF). The positioning of Insert 2 of SP15- like DNAPs is similar to that of H5-H5’ homodimer of Mpox and partially overlaps the *γ*B subunit of mitochondria, while Insert 1 of SP15-like DNAPs and A22 in Mpox DNAP holoenzyme is localized on the opposite side of the DNA.

On the lagging strand, the UDG domain precedes the replicative DNA polymerase, engaging with the 5′-end single-stranded DNA (ssDNA) template. Strikingly, the positioning of the UDG domain is similar to the 5’-3’ exo/endonuclease domain (Fen1-like) of A-Family DNA polymerase in the *Escherichia coli* replication machinery^45^, suggesting also a possible strand displacement function for this domain (**Suppl. Fig. 8**).

Many phage genomes coming from metagenomics project contain a UDG protein in trans of the DNAP^59^ and the present structures allow to predict how it associates with the DNAP, at least if it is an A-Family DNAP. Strikingly, the UDG domain occupies the same position in the replication complex of mpox virus (which involves a B-family DNA polymerase). Also, Insert 2 structurally resembles the H5-H5’ hetero-tetramer in Monkey-pox virus or the UL42 protein in herpes virus (**Fig. 8c**) that act as processivity factors in these human viruses^44^, while Insert1 occupies the same location than A22 subunit in Mpox complex. Altogether, Inserts 1 and 2 occupy analogous positions to PCNA (β clamp in eukaryotes) in archaea and bacteria, suggesting again a conserved role in stabilizing the DNA replication machinery.

Beyond the viral world, two A-Family DNAPs stand out in eukaryotes: the polymerase theta, involved in DNA repair, and the replicative polymerase gamma (Pol γ) in mitochondria, which naturally associates closely with the γ2 accessory subunit that forms a homodimer, forming a heterotrimer^60–62^. Intriguingly, there is one nuclear-encoded mitochondrial glycosylase called OGG1 in eukaryotes, involved in the excision of the oxidized base 8-oxo-G, and the work presented here suggests that it may associate directly with the A-family Pol γ in a way that is illustrated in **Fig. 8c**. Further work will be needed to characterize the interaction interface at the molecular level.

### Limitations and opportunities

Currently, only two of SP15-like DNAPs are confirmed to come from phages with heavily T-modified DNA, especially through glycosylation and a long linker (YerA41 and SP-15). Genome analysis further suggests that RP13, EP_H11 and EcoM_IME92 are also T-modified due to the presence of the aGPT-Pplase gene. Also, all members have genes for dCTP deaminase, a dUTP phosphorylase and a thymidylate synthase homologue. It would be interesting to study if this is a general property of all the phages in this group. Notably, this modification of their DNA is important not only for their resistance to the host endonucleases, but also for their packaging.

We note that a psi-blast search using YerA41 DNAP X retrieves not only the corresponding gene of SP15-like phages but also numerous Nucleo-Cytoplasmic Large DNA viruses that all have a dUTPase, a UDG protein and a DNA ligase. It would be interesting to check if the DNAP X interacts with the replicative DNAP, which in this case is a B-family DNA polymerase.

Here we describe two members of this family as multifunctional replicative polymerases and infer that they serve as a platform to recruit a X-family DNAP and DNA ligase to repair dU bases, with a stable interaction between the replicative A-family and the repair X-family DNAPs. This provides a model for an integrated BER-like pathway in these phages (**Fig. 7e**). While no strong interaction was detected between DNAP X and DNA ligase (**Fig. 7b**), we note the recent report of a fused DNAP X and DNA ligase in certain giant dsDNA viruses, such as PgV-16T, which likely facilitates gap repair through a coordinated exchange of catalytic domains depending on the substrate^63^. Further structural characterization will help elucidate how A-family DNAPs function as scaffolds for BER components.

In the future, the ability to synthesize sugar-modified T-containing DNA templates similar to SP-15 may facilitate structural studies advancing the molecular understanding of how SP15-like DNAPs accommodate various types of hypermodified DNA substrates, and perhaps how to engineer them to make them more general in front of yet unknown T-modifications.

## Materials and Methods

### Blast search, sequence analysis, gene neighborhood analysis, structure prediction and phylogeny

YerA_144, the A-family DNA polymerase from *Yersinia ruckeri* phage, consists of 1,306 amino acid residues. Sequence similarity analysis was conducted using BLAST^64^ (version 2.16.0), retaining only DNA polymerase sequences of at least 1,000 amino acids.

A multiple sequence alignment (MSA) was performed using Clustal Omega on the EMBL web server ^65^, and manually checked for the presence of known important motifs both in polymerase and UDG domains, refined and visualized using Espript^66,67^ (see **Suppl. Fig. 1**). Fourteen DNA polymerase sequences were selected for further analysis, and gene neighborhood analysis was conducted for each viral source.

DNA polymerases from the X-family and DNA ligases were identified where present in the respective viral genomes. Genes encoding A-family DNA polymerases, X-family DNA polymerases, and DNA ligases from each virus were inventoried and summarized in **Suppl. Table 2**, which includes their respective gene names.

The DNA polymerase sequences homologous to (*gp144*) were pooled with the dataset of 8,136 sequences from the study by Czernecki^31^ and processed using the CLANS web server from the MPI Bioinformatics Toolkit^68^. Clustering simulations were carried out using the JAVA version of CLANS^32^, running for over 8,000 iterations to ensure robust clustering results.

AlphaFold3 ^69^ was used to produce models from UDG polymerases, including SPO1-like polymerases, N4 polymerases, and SP15-like polymerases.

### Molecular cloning and protein expression

The wild-type synthetic gene *gp144* was kindly provided by Dr. M. Skurnik and used as described in a previous study^8^. The wild-type synthetic genes for DNA polymerase β from *Yersinia ruckeri* phage (*gp127*), DNA ligase from *Yersinia ruckeri* phage (*gp158*), and DNA polymerase I from *Thermus thermophilus* phage (*gp158*) were optimized for expression in *Escherichia coli* (**Suppl. Table 3**) and synthesized by GeneScript. These genes were cloned into a modified pRSF1-Duet expression vector, which includes an N-terminal 14-histidine tag and a TEV protease cleavage site. Exonuclease-deficient mutants (Exo-) of the DNA polymerases from YerA and phiLo phages were subcloned for further studies. The point mutations (D263A-E265A in phiLo_158) were introduced via PCR using primers designed from Eurogentec (Suppl. Table S3) and Q5 Hot Start High-Fidelity DNA Polymerase (NEB, M0493S). PCR products were subsequently treated with a Kinase, Ligase, and DpnI (KLD) kit (NEB, M0554S) before transformation into *E. coli* DH5⍺ cells with the newly constructed exo-plasmids and were verified by Sanger sequencing by Eurofins.

Escherichia coli BL21 Star (DE3) cells (Invitrogen) were transformed with the engineered plasmids. Bacteria were cultivated at 37^◦^C in LB medium with kanamycin resistance selection and induced at an optical density (OD) of 0.6–0.8 with 0.5 mM isopropyl-β-D-thiogalactopyranoside (IPTG). After incubation overnight at 20°C for YerA_127, YerA_158, phiLo_158 and 15°C for YerA_144, cells were harvested and homogenized in suspension buffer (50 mM HEPES pH 8.0, 300 mM NaCl, 10 mM imidazole).

### Protein purifications

After sonication and centrifugation of bacterial debris, corresponding lysate supernatants were supplemented with Benzonase (Sigma-Aldrich) and protease inhibitors (Thermo Fisher Scientific), 1 µl and one tablet per 50 ml, respectively. The proteins of interest were isolated by purification of the lysates on a HisTrap column (suspension buffer as washing buffer, 500 mM imidazole in elution buffer). Collected proteins were diluted to 50 mM NaCl and repurified on a HiTrap Heparin column with an elution at 1 M NaCl. Both purification columns were from Cytiva. Protein purity was assessed on a sodium dodecyl sulphate–polyacrylamide gel electrophoresis (SDS–PAGE) 4–15% gel (BioRad) with a molecular weight ladder (Precision Plus Protein, Biorad) as control. The enzymes were concentrated with Amicon Ultra 30k MWCO centrifugal filters (Merck), flash-frozen in liquid nitrogen and stored directly at −80°C, with no glycerol added.

### DNA substrates constructs

DNA oligomers (Eurofins) were dissolved in Tris-EDTA buffer pH 8 at 100 µM. Complementary oligonucleotides were mixed at an equimolar ratio in 25 mM HEPES–NaOH (pH 7.5), 150 mM sodium acetate, 2 mM magnesium acetate and annealed by gradual cooling from 90 °C to room temperature on night.

Oligomer sequences were as follows:

Pair 1: oSM0072 5’-CACAGACGTAGCAGCTGGAACCTCCGTA-3’ oSM0073 5’-AAAA[U]CCCCGCTACGGAGGTTCCAGCTGCTACGTCTGTG-3’
Pair 2: oSM0072 5’-CACAGACGTAGCAGCTGGAACCTCCGTA-3’ oSM0074 5’-AAAATCCCCCCTACGGAGGTTCCAGCTGCTACGTCTGTG-3’

### Preparation of complexes for cryo EM

To assemble the first complex, 20 μM exo-YerA41_144 was mixed with the duplex osM0072/oSM0074 at a 1:1.2 molar ratio in a buffer containing 20 mM HEPES pH 8, 50 mM NaCl, 5 mM DTT, and 5 mM MgCl_2_ and incubated for 5 minutes at 4° prior to grids preparation.

A second complex of exo– phiLo_158 was prepared as mentioned above.

A third complex of exo-phiLo_158 was prepared as mentioned above but was mixed with the duplex oSM0072 and 0SM0073. The complexes were applied to negatively glow-discharged 300 mesh – 2/1 holey-gold grids (Quantifoil). Grids were blotted with ash-free Whatman® Grade 540 filter paper in a Vitrobot Mark IV (ThermoFisher Scientific) for 4 s at 4 °C and 95–100% humidity, then vitrified in liquid ethane. The sample quality and distribution were assessed using Glacios Transmission Electron Microscope equipped with a Falcon 4 direct electron detector.

### Cryo EM data collections & data processing

Datas from each complex were collected at the Nanoimaging facility for CryoEM at Pasteur Institute using a Glacios transmission electron microscope (ThermoFisher Scientific), operated at 200 kV and equipped with a Bioquantum Energy Filter with a 10 eV slit width. Movies were collected using a Falcon direct electron detector in super-resolution mode with a magnification of 205,000, corresponding to a pixel size of 0.57 Å. A dose rate of 15 e^−^/s/physical pixel resulted in a total electron dose of 40 e^−^/Å^2^, which was applied over 40 frames. Data was collected in EPU software with defocus values ranging from −1 to −3 µm.

The movie stacks collected for were processed in CryoSPARC^1^. The super-resolution movies were frame-aligned, motion-corrected, gain-normalized, dose-weighted, and binned twice with the patch motion correction module. Contrast transfer function (CTF) values were estimated using the patch CTF (CryoSPARC)^71^. Micrographs with ice, ethane contamination, and/or poor CTF fit resolution were discarded (below 5 Å). A circular blob picker with dimensions of 80–130 Å was used to pick particles. Local resolution plots were obtained in CryoSPARC. The reported resolutions of the CryoEM maps are based on FSC 0.143 criterion.

An additional dataset of DNAP YERA41 in complex with DNA was collected on a Titan Krios G4, equipped with a cold FEG, a Selectris X energy filter, a Falcon 4i, and a Volta Phase Plate. Data collection was performed at CM02 of the ESRF CRG line (Grenoble). Movies were collected using a Falcon 4i direct electron detector in super-resolution mode with a magnification of 205,000, corresponding to a pixel size of 0.57 Å. Data were acquired with an applied tilt of 0° and 30° to address the preferred orientation bias of the sample and a defocus range from −1 to −2.4 µm. A dose rate of 15 e⁻/s/physical pixel resulted in a total electron dose of 40 e⁻/Å², distributed over 40 frames. A similar workflow was applied to this last dataset. Workflows for image processing of complex are shown in **Suppl. Fig. 2-3-4**, respectively.

### Molecular modelling

The initial structural model was generated using RosettaFold and AlphaFold, two structure prediction tools that leverage deep learning methods to predict accurate atomic models based on sequence data. The resulting models provided a reliable starting point for subsequent refinement. These initial predictions were then rigid-body fitted into the cryo-EM density map using tools in ChimeraX^72^ to optimize the global placement. To improve the agreement between the model and the experimental map, molecular modeling and flexible fitting were carried out using real-space refinement in software such as Phenix ^73^ and COOT ^74^. This iterative process ensured that the structure conformed to the high-resolution cryo-EM map with correct stereochemistry, resolving ambiguities and accommodating local conformational adjustments. Ultimately, this workflow yielded an accurate and high-quality atomic model that fit well into the cryo-EM density, enabling further structural analysis.

### Activity assay

Polymerase activity tests were performed in 20 mM Tris-HCl pH 8.0, 50 mM NaCl, 5 mM MgCl_2_, 10 mM DTT. Reaction solutions contained 1 µM of Cy5-templating oligonucleotides, 1 µM of 5’FAM- labeled DNA primer (all oligonucleotides were purchased from Eurofins and are listed in **Supp. Table 2**), 1 mM of each dNTPs, 1 µM of DNA polymerases, one unit of APE1 (NEB), one unit of UDG protein (NEB).

Bypass of THF site were performed in the same buffer activity. Reactions contained 1 µM of DNA template oligonucleotide and 1 µM of 5’FAM-labeled primer, 1 µM of nucleotides mix or of single nucleotides.

Apurinic endonuclease activity tests were performed using 1 µM template 5’-Cy5-labeled DNA strand containing a dU base or THF site or a gap 5’-labeled with Cy5, 1 µM 5’-FAM-labeled primer, 1 mM of dNTPs mix, 1 µM of DNA polymerase X from YerA41 and finally DNA ligase from YerA41, pre incubated with 1 mM of NAD+.

## Acknowledgements

We would like to thank the staff of the Nanoimaging Core Facility at Institut Pasteur for providing access to the cryo-EM instruments for collecting the cryo-EM datasets. We also thank the ESRF CM02 beamline staff for assisting in the collection of the final dataset of DNA polymerase I YerA41 in tilt mode. We thank also Milena Milovanovic for doing the pre-characterization of phiLo DNAP activity. We would like to express our gratitude to Dr. Michael Skurnik for providing the DNA polymerase I YerA41 plasmid and for regular scientific discussions. We thank Dr. Peter R. Weigele and Dr. Laurence Ettwiller from New England Biolabs, USA, for exchanging scientific information and encouragement.

## Financial support

We acknowledge partial financial support from the ANR BreakDance grant as well as the PEPR grant MoleculRxiv. ADMB lab is also supported by Institut Pasteur and CNRS.

## Deposited data

The structural models have been deposited in the PBD with codes 9QHS, 9QHT, 9QHU, 9QLH and 9QLI. The Carin-5 genome has been deposited in Genbank and is available at https://www.ncbi.nlm.nih.gov/nuccore/PV155638

## Author contributions

S.M. and M.D. conceived the experiments and analyzed the data. S.M. performed molecular cloning, protein expression, and purification; prepared the cryo-EM grids; collected and analyzed the data; reconstructed the cryo-EM maps and built the models. S.M. and S.L.S. independently performed activity assays, conducting at least three replicates. R.S. assisted in designing the activity assays and in the bioinformatics analysis and contributed data with the Carin5 genome. S.M. and M.D. co-wrote the paper.

## Competing interests

The authors declare no competing interests.

## Notes

### Competing Interest Statement

The authors have declared no competing interest.

## References

1. Kornberg, A. DNA Replication. New York: W.H. Freeman and Company (1980).

2. Warren, R. A. Modified bases in bacteriophage DNAs. Annu. Rev. Microbiol. 34, 137–158 (1980).

3. H.Gommers-Ampt, J. & Borst, P. Hypermodified bases in DNA. FASEB J. 9, 1034–1042 (1995).

4. Drozdz, M., Piekarowicz, A., Bujnicki, J. M. & Radlinska, M. Novel non-specific DNA adenine methyltransferases. Nucleic Acids Res. 40, 2119–2130 (2012).

5. Weigele, P. & Raleigh, E. A. Biosynthesis and Function of Modified Bases in Bacteria and Their Viruses. Chem. Rev. 116, 12655–12687 (2016).

6. Hutinet, G. et al. 7-Deazaguanine modifications protect phage DNA from host restriction systems. Nat. Commun. 10, 5442 (2019).

7. Lee, Y.-J. et al. Identification and biosynthesis of thymidine hypermodifications in the genomic DNA of widespread bacterial viruses. Proc. Natl. Acad. Sci. U. S. A. 115, E3116–E3125 (2018).

8. Gomez-Raya-Vilanova, M. V. et al. The DNA polymerase of bacteriophage YerA41 replicates its T-modified DNA in a primer-independent manner. Nucleic Acids Res. 50, 3985–3997 (2022).

9. Wang, S. et al. Landscape of New Nuclease-Containing Antiphage Systems in Escherichia coli and the Counterdefense Roles of Bacteriophage T4 Genome Modifications. J. Virol. 97, e00599–23 (2023).

10. Krupovic, M., Dolja, V. V. & Koonin, E. V. Origin of viruses: primordial replicators recruiting capsids from hosts. Nat. Rev. Microbiol. 17, 449–458 (2019).

11. Sleiman, D. et al. A third purine biosynthetic pathway encoded by aminoadenine-based viral DNA genomes. Science 372, 516–520 (2021).

12. Czernecki, D. et al. How cyanophage S-2L rejects adenine and incorporates 2-aminoadenine to saturate hydrogen bonding in its DNA. Nat. Commun. 12, 2420 (2021).

13. Zhou, Y. et al. A widespread pathway for substitution of adenine by diaminopurine in phage genomes. Science 372, 512–516 (2021).

14. Pezo, V. et al. Noncanonical DNA polymerization by aminoadenine-based siphoviruses. Science 372, 520–524 (2021).

15. Kawasaki, T., Endo, H., Ogata, H., Chatchawankanphanich, O. & Yamada, T. The complete genomic sequence of the novel myovirus RP13 infecting Ralstonia solanacearum, the causative agent of bacterial wilt. Arch. Virol. 166, 651–654 (2021).

16. Leskinen, K. et al. YerA41, a Yersinia ruckeri Bacteriophage: Determination of a Non-Sequencable DNA Bacteriophage Genome via RNA-Sequencing. Viruses 12, 620 (2020).

17. Hossain, A. A. et al. DNA glycosylases provide antiviral defence in prokaryotes. Nature 629, 410–416 (2024).

18. Kornberg, A. DNA replication. J. Biol. Chem. 263, 1–4 (1988).

19. Yutin, N. et al. Analysis of metagenome-assembled viral genomes from the human gut reveals diverse putative CrAss-like phages with unique genomic features. Nat. Commun. 12, 1044 (2021).

20. Katz, G. E., Price, A. R. & Pomerantz, M. J. Bacteriophage PBS2-induced inhibition of uracil-containing DNA degradation. J VIROL **Vol** 20, 535–538 (1976).

21. Serrano-Heras, G., Bravo, A. & Salas, M. Phage φ29 protein p56 prevents viral DNA replication impairment caused by uracil excision activity of uracil-DNA glycosylase. Proc. Natl. Acad. Sci. 105, 19044–19049 (2008).

22. Longo, M. C., Berninger, M. S. & Hartley, J. L. Use of uracil DNA glycosylase to control carry-over contamination in polymerase chain reactions. Gene 93, 125–128 (1990).

23. Schormann, N., Ricciardi, R. & Chattopadhyay, D. Uracil-DNA glycosylases—Structural and functional perspectives on an essential family of DNA repair enzymes. Protein Sci. Publ. Protein Soc. 23, 1667–1685 (2014).

24. Friedberg, E. C. DNA damage and repair. Nature 421, 436–440 (2003).

25. Ahn, W.-C. et al. Covalent binding of uracil DNA glycosylase UdgX to abasic DNA upon uracil excision. Nat. Chem. Biol. 15, 607–614 (2019).

26. Shiraishi, M., Ishino, S., Heffernan, M., Cann, I. & Ishino, Y. The mesophilic archaeon Methanosarcina acetivorans counteracts uracil in DNA with multiple enzymes: EndoQ, ExoIII, and UDG. Sci. Rep. 8, 15791 (2018).

27. Bogani, F., Chua, C. N. & Boehmer, P. E. Reconstitution of Uracil DNA Glycosylase-initiated Base Excision Repair in Herpes Simplex Virus-1*. J. Biol. Chem. 284, 16784–16790 (2009).

28. Xu, Y. et al. Cryo-EM structures of human monkeypox viral replication complexes with and without DNA duplex. Cell Res. 33, 479–482 (2023).

29. Peng, Q. et al. Structure of monkeypox virus DNA polymerase holoenzyme. Science 379, 100–105 (2023).

30. Li, Y., Shen, Y., Hu, Z. & Yan, R. Structural basis for the assembly of the DNA polymerase holoenzyme from a monkeypox virus variant. Sci. Adv. 9, eadg2331 (2023).

31. Czernecki, D., Nourisson, A., Legrand, P. & Delarue, M. Reclassification of family A DNA polymerases reveals novel functional subfamilies and distinctive structural features. Nucleic Acids Res. 51, 4488–4507 (2023).

32. Frickey, T. & Lupas, A. CLANS: a Java application for visualizing protein families based on pairwise similarity. Bioinformatics 20, 3702–3704 (2004).

33. Ehrlich, M. & Ehrlich, K. C. A novel, highly modified, bacteriophage DNA in which thymine is partly replaced by a phosphoglucuronate moiety covalently bound to 5-(4‘,5‘- dihydroxypentyl)uracil. J. Biol. Chem. 256, 9966–9972 (1981).

34. Iyer, L. M., Zhang, D., Maxwell Burroughs, A. & Aravind, L. Computational identification of novel biochemical systems involved in oxidation, glycosylation and other complex modifications of bases in DNA. Nucleic Acids Res. 41, 7635–7655 (2013).

35. Lee, Y.-J. et al. Pathways of thymidine hypermodification. Nucleic Acids Res. 50, 3001–3017 (2022).

36. Kropinski, A. M. et al. The Sequence of Two Bacteriophages with Hypermodified Bases Reveals Novel Phage-Host Interactions. Viruses 10, 217 (2018).

37. Stewart, C. R. et al. The genome of Bacillus subtilis bacteriophage SPO1. J. Mol. Biol. 388, 48–70 (2009).

38. Jumper, J. et al. Highly accurate protein structure prediction with AlphaFold. Nature 596, 583–589 (2021).

39. Mukherjee, S. & Zhang, Y. MM-align: a quick algorithm for aligning multiple-chain protein complex structures using iterative dynamic programming. Nucleic Acids Res. 37, e83–e83 (2009).

40. Steitz, T. A. DNA polymerases: structural diversity and common mechanisms. J. Biol. Chem. 274, 17395–17398 (1999).

41. Doublié, S., Sawaya, M. R. & Ellenberger, T. An open and closed case for all polymerases. Struct. Lond. Engl. 1993 7, R31–35 (1999).

42. Rothwell, P. J. & Waksman, G. Structure and mechanism of DNA polymerases. Adv. Protein Chem. 71, 401–440 (2005).

43. Akabayov, B. et al. Conformational dynamics of bacteriophage T7 DNA polymerase and its processivity factor, Escherichia coli thioredoxin. Proc. Natl. Acad. Sci. 107, 15033–15038 (2010).

44. Wang, X., Ma, L., Li, N. & Gao, N. Structural insights into the assembly and mechanism of mpox virus DNA polymerase complex F8-A22-E4-H5. Mol. Cell 83, 4398–4412.e4 (2023).

45. Botto, M. M., Borsellini, A. & Lamers, M. H. A four-point molecular handover during Okazaki maturation. Nat. Struct. Mol. Biol. 30, 1505–1515 (2023).

46. Stuart, D. T., Upton, C., Higman, M. A., Niles, E. G. & McFadden, G. A poxvirus-encoded uracil DNA glycosylase is essential for virus viability. J. Virol. 67, 2503–2512 (1993).

47. Hogg, M., Wallace, S. S. & Doublié, S. Crystallographic snapshots of a replicative DNA polymerase encountering an abasic site. EMBO J. 23, 1483–1493 (2004).

48. Obeid, S. et al. Replication through an abasic DNA lesion: structural basis for adenine selectivity. EMBO J. 29, 1738–1747 (2010).

49. Hogg, M., Seki, M., Wood, R. D., Doublié, S. & Wallace, S. S. Lesion bypass activity of DNA polymerase θ (POLQ) is an intrinsic property of the pol domain and depends on unique sequence inserts. J. Mol. Biol. 405, 642–652 (2011).

50. Bogani, F. & Boehmer, P. E. The replicative DNA polymerase of herpes simplex virus 1 exhibits apurinic/apyrimidinic and 5′-deoxyribose phosphate lyase activities. Proc. Natl. Acad. Sci. 105, 11709–11714 (2008).

51. García-Escudero, R., García-Díaz, M., Salas, M. L., Blanco, L. & Salas, J. DNA Polymerase X of African Swine Fever Virus: Insertion Fidelity on Gapped DNA substrates and AP lyase Activity Support a Role in Base Excision Repair of Viral DNA. J. Mol. Biol. 326, 1403–1412 (2003).

52. Redrejo-Rodríguez, M., García-Escudero, R., Yáñez-Muñoz, R. J., Salas, M. L. & Salas, J. African Swine Fever Virus Protein pE296R Is a DNA Repair Apurinic/Apyrimidinic Endonuclease Required for Virus Growth in Swine Macrophages. J. Virol. 80, 4847–4857 (2006).

53. Baños, B., Villar, L., Salas, M. & de Vega, M. Intrinsic apurinic/apyrimidinic (AP) endonuclease activity enables Bacillus subtilis DNA polymerase X to recognize, incise, and further repair abasic sites. Proc. Natl. Acad. Sci. U. S. A. 107, 19219–19224 (2010).

54. Pérez-Lago, L. et al. Characterization of Bacillus subtilis uracil-DNA glycosylase and its inhibition by phage φ29 protein p56. Mol. Microbiol. 80, 1657–1666 (2011).

55. Obeid, S. et al. Replication through an abasic DNA lesion: structural basis for adenine selectivity. EMBO J. 29, 1738–1747 (2010).

56. Gao, Y. et al. Structures and operating principles of the replisome. Science 363, eaav7003 (2019).

57. Wallen, J. R. et al. Hybrid Methods Reveal Multiple Flexibly Linked DNA Polymerases within the Bacteriophage T7 Replisome. Structure 25, 157–166 (2017).

58. Kulczyk, A. W., Moeller, A., Meyer, P., Sliz, P. & Richardson, C. C. Cryo-EM structure of the replisome reveals multiple interactions coordinating DNA synthesis. Proc. Natl. Acad. Sci. 114, E1848–E1856 (2017).

59. Yutin, N. et al. DNA polymerase swapping in Caudoviricetes bacteriophages. Virol. J. 21, 200 (2024).

60. Lim, S. E., Longley, M. J. & Copeland, W. C. The mitochondrial p55 accessory subunit of human DNA polymerase gamma enhances DNA binding, promotes processive DNA synthesis, and confers N-ethylmaleimide resistance. J. Biol. Chem. 274, 38197–38203 (1999).

61. Wojtaszek, J. L. et al. Structure-specific roles for PolG2–DNA complexes in maintenance and replication of mitochondrial DNA. Nucleic Acids Res. 51, 9716–9732 (2023).

62. Johnson, A. A., Tsai, Y. c, Graves, S. W. & Johnson, K. A. Human mitochondrial DNA polymerase holoenzyme: reconstitution and characterization. Biochemistry 39, 1702–1708 (2000).

63. Fernández-García, J. L., de Ory, A., Brussaard, C. P. D. & de Vega, M. Phaeocystis globosa Virus DNA Polymerase X: a “Swiss Army knife”, Multifunctional DNA polymerase-lyase-ligase for Base Excision Repair. Sci. Rep. 7, 6907 (2017).

64. Altschul, S. F. et al. Gapped BLAST and PSI-BLAST: a new generation of protein database search programs. Nucleic Acids Res. 25, 3389–3402 (1997).

65. Madeira, F. et al. The EMBL-EBI Job Dispatcher sequence analysis tools framework in 2024. Nucleic Acids Res. 52, W521–W525 (2024).

66. Gouet, P., Courcelle, E., Stuart, D. I. & Metoz, F. ESPript: analysis of multiple sequence alignments in PostScript. Bioinformatics 15, 305–308 (1999).

67. Robert, X. & Gouet, P. Deciphering key features in protein structures with the new ENDscript server. Nucleic Acids Res. 42, W320–W324 (2014).

68. Gabler, F., et al. Protein Sequence Analysis Using the MPI Bioinformatics Toolkit. Curr. Protoc. Bioinforma. 72, e108 (2020).

69. Abramson, J. et al. Accurate structure prediction of biomolecular interactions with AlphaFold 3. Nature 630, 493–500 (2024).

70. Rubinstein, J. L. & Brubaker, M. A. Alignment of cryo-EM movies of individual particles by optimization of image translations. J. Struct. Biol. 192, 188–195 (2015).

71. Rohou, A. & Grigorieff, N. CTFFIND4: Fast and accurate defocus estimation from electron micrographs. J. Struct. Biol. 192, 216–221 (2015).

72. Meng, E. C. et al. UCSF ChimeraX: Tools for structure building and analysis. Protein Sci. Publ. Protein Soc. 32, e4792 (2023).

73. Afonine, P. V. et al. Real-space refinement in PHENIX for cryo-EM and crystallography. Acta Crystallogr. Sect. Struct. Biol. 74, 531–544 (2018).

74. Emsley, P., Lohkamp, B., Scott, W. G. & Cowtan, K. Features and development of *Coot*. *Acta Crystallogr*. D Biol. Crystallogr. 66, 486–501 (2010).

