## Supplementary Material for "Structural Basis for DNA Replication and Uracil Repair in Phage A-Family DNA Polymerases"

-Supplementary Figure 1
-Supplementary Figure 2
-Supplementary Figure 3
-Supplementary Figure 4
-Supplementary Figure 5
-Supplementary Figure 6
-Supplementary Figure 7
-Supplementary Figure 8
-Supplementary Table 1
-Supplementary Table 2
-Supplementary Table 3
-References

```
                                                                   UDG A
                  1                                         10
RP13/1-1017       MHI.......................................VRSTNEKS.K..VLVIGD
YerA41/1-1218     .................................................M.S..TVLIGY
phiLo/1-1134      MRETEREVVEVYRVLPIAREDKCNSCPLKGNFP.....HSVGYEFYGEPPYE..YLLVGE
RHph_Y65/1-1067   MSLEIESIINKVDLTSDVR..VCHMCPLAEKFGGCDNLGRVSFEIVGQNP.K..VLLVGE
EP_H11/1-1049     MFL.......................................NHNLVGEV.D..YVIIGD
SP-15/1-1140      MA..........DTSTSKEI......................FPIGTLNP.T..VYIVGE
Carin5/1-1216     .................................................M.SQKIALIGY
LIS04/1-1220      MANYAIDGPIGVD...................................NP.E..VYIVGE

                  20        30         40              50
RP13/1-1017       SVITPDMNPVVPFDSEAGNSILKPLMMV......AGVYPH..YMDIKKD...........
YerA41/1-1218     ....VNIRP.....GKGGKLINQDVWKYFEKYVPAHFEDV..DLLLF.............
phiLo/1-1134      APGNDEISWGRPF.IGKSGQLLRQALSE......VGISSYI.............LTNVSM
RHph_Y65/1-1067   APAAEDLKSERPF.SSQSGSFLRMAMAE......IGINDYA.............LANVIM
EP_H11/1-1049     E...RENNDAPPFAASNAGQTLDVALIT......AGVHNKVGLVEINRDGTFDELSNIDE
SP-15/1-1140      SPGAI.........MGQAGTYLQKITRS......LGL.......................
Carin5/1-1216     ....VNIKK.....NQYGKVVNKDIWQYFSAYVEEHFENV..DLLLF.............
LIS04/1-1220      AANSI.........LSNSGQYLQKVTQS......LGLTPF....................

                   60                                                        70
RP13/1-1017       ..D..YD.......QEMTPEV......................................L
YerA41/1-1218     ..............NPQTLPN......................................L
phiLo/1-1134      CIPVVPGSKVAPPPKEAASYC.....................................SS
RHph_Y65/1-1067   CQP..KSVNIYTPTPEESAHC......................................K
EP_H11/1-1049     IRD..KD.......GLNFYEL......................................C
SP-15/1-1140      .................................DTEINLRMFSISRKFGEVDASYMTQSS
Carin5/1-1216     ..............NNDTLPS......................................L
LIS04/1-1220      .....KNLRLLTLTREWTSDSSTLDSTSMDVIH...........................

                          80         90
RP13/1-1017       AEVFAYISANDIKQVITLGRWSTQSL..................................
YerA41/1-1218     SDMIEDLNKYD..NIIVTGREPLKYLGL................................
phiLo/1-1134      SYLVPFIQRVRPKKVVLLGRTAIEFV..................................
RHph_Y65/1-1067   AHIQRFVEKYKPETVILLGRGAYEVI..................................
EP_H11/1-1049     AK.......HNVKHVMTLGRQLAATI..................................
SP-15/1-1140      RFVLADLAKTKPKVIIGLGATPAGLL..................................
Carin5/1-1216     AGMTAKLNQYD..SLIVAGREPLKFLDLAAVESDEEDVVEDEESEEDDSDDSDEDDDDDA
LIS04/1-1220      .....DIARTKPKVVVGLGAVPAKVF..................................

                                                                         UDG B
                                100                     110        120        130
RP13/1-1017       ...........F..P.SAI...........KKQTMAWLTSIPMFEPMELQGIKLGVMYSP
YerA41/1-1218     .EDAEIDRHSQFSNK.SDN...........KIEVMRYFGLINNNPALKFNKIH..VMYSL
phiLo/1-1134      ...........L..K.ALLGERFKEGGISDYKKT.........YFGVEVDGIHYFPLNHP
RHph_Y65/1-1067   ...........L..PESYF...........AGQS...VTNVTKKKAIEIDGVRYGAAFHP
EP_H11/1-1049     ...........S..P.GYT...........HKKTLDKLLKFPLGNIPEY.GFKLAVVMHH
SP-15/1-1140      ...........T..G........SMGRVSDIRGNVEKIT.......VGDHECYFITTFHP
Carin5/1-1216     DENEEVNK.....NN.SSS...........KLDDLRFHGIVNNNPELKFEKID..VIFSL
LIS04/1-1220      ...........T..G........IDGKVSDLRGNISEIT.......VEGHQCYFITTFHP

                              140        150
RP13/1-1017       MQVLKS........NDPQAAEEWSTRLEKLFTVT.......................D..
YerA41/1-1218     MMYIAK........DKINVFH.........FDLL.......................NRT
phiLo/1-1134      SFVLRRGG......VESQEYKTFVGELQSVVEFSHVQVEIPSDPVSFSHLFSTGKIPV..
RHph_Y65/1-1067   SYIFRNGG......KESPQYIDFLNRLRRLTQNS.......................V..
EP_H11/1-1049     AQVERNLD......SDPGLLDSWIDRFKFLVFGQ.......................K..
SP-15/1-1140      GYILNNAGNTAIMKT.......FLGDFRKAIAAS..................EGKVDA..
Carin5/1-1216     IYYISK........DLINVFH.........YDML.......................NRT
LIS04/1-1220      NYILNN........TSTSNAKLFVGDFRKAVAAA..................SGKVNT..

                                                                   EXO I
                            160        170                180        190        200
RP13/1-1017       ..........EVVVNVKHIETEDEFI......ELLNHLDSFPVGRQRGLDYETNAVDAFN
YerA41/1-1218     GKNNLVPPMVQIDPKFHH....ISEL......ESIYHSK.WQPDEHKGFDFETR...NFP
phiLo/1-1134      ..........SPKGRL.....NYHVFDLDSYEEGFSL...IDDLDEVSFDVE.TSKGETS
RHph_Y65/1-1067   ..........DAEFIV.....PFKIVNLDQLDETLEE...FDVYKELGFDYEGTSLDPYD
EP_H11/1-1049     ..........ESEYNV.....EWEFVDWNGLQEFLNDIMWSGYKGYLGLDYESNAADLFS
SP-15/1-1140      ..........NSDKDL.....RIALTAQEFETFLKNE...YTADGETSYDIETNGADELS
Carin5/1-1216     GTNHAIPEMGDIDPKYHH....ISEL......KSIFDSQ.YLPDESIGFDFETR...MFP
LIS04/1-1220      ..........DSEKDL.....RVALTADDFINFCEQN...YTSNGETSYDIETNGKDVLS
```

```
                          210       220       230              240         250
RP13/1-1017       VDHYITMIGLSVLISPTEA.NAWYWVPPRKM...SD...RVLEAFTNFLVKW....KNDI
YerA41/1-1218     FEPRFRLLGFSLA.NDTYQ.AYFYFSRDDDK...YA...QEMEQYRSVMVKIFLEDAKKL
phiLo/1-1134      VDRDLVLVGLGNH.....EVQVAFDL.REGS...EENAPRFLQRLREYLDKP....GRKV
RHph_Y65/1-1067   VNYQVLGLGMSSM.....DKAVYIDFHREPT...QE...E.KNKISDFIRKK....Q..M
EP_H11/1-1049     VDFRITGFSIAKKIGDKEAKSVWYNVTGKET...PE...Q.LMFLRDFLVKF....QELI
SP-15/1-1140      NKAKVVGFSIAPD....EVKGIYVIFEALEYKMPEEDVERCKTLLRDFLKER....R..V
Carin5/1-1216     FWSNFRILGFSLV.SNTYQ.AYFYFSRDDEK...YI...QEFNEYRQDLLDIFNSNHKRF
LIS04/1-1220      DKARVVGFSVSPD....DKVGTYVVFEALEYQMPESDVERCKEYLRNFLSER....H..I

                     EXO II
                           260          270      280            290      300
RP13/1-1017       WAYNVTFEMKVT....WHAT.GKYVEMQDALVLATCLA......KRGSLKDNVRAEFGAG
YerA41/1-1218     HTYNNKFENNAV....LDQL.RERIFFEDSRIYTQMLK......TRGSLKFSANLFLGVP
phiLo/1-1134      IVYNQSFEAGVY....LQLY.GEFRSFVDVFSMTKAFGLQTELYQKIGLKKTASVIFGVP
RHph_Y65/1-1067   WVYNVNYETKVT....WSKLSNDLIRFHDAMTLVKIDC......SPGSLKDNARKYLNAE
EP_H11/1-1049     VAFNCPYEIKVT....WHWL.DVFCKFIDAIVTLTMHG......ERSSLKDAARRYLHAD
SP-15/1-1140      IIHNQMYERPYTLNENWLGY.DLHDNIDDTLVMSRLML...GGKTGAGLKDQCRLNLGYT
Carin5/1-1216     WAYNNKFENNVV....FDQF.KTPIFFEDARIFTQILK......TRGSLKFSANLFLGVP
LIS04/1-1220      VVHNQLYERPYTLNENWLGY.ELHDDMDDTLVMSRLML...GGNTGAGLKDQCRLNLGYT

                            310       320
RP13/1-1017       LWEAPVHAFKAKSGELFKIVMAAKP..................I................
YerA41/1-1218     VWNEGINTFMKNYKEFVAWMKDYREL..........SMIEESVI................
phiLo/1-1134      EWKDRVLDYVRLAEDMSENLEDWR....................................
RHph_Y65/1-1067   IWEEIVGNLTTLFYDIFQFMKKIE....................................
EP_H11/1-1049     FWEGSVHEWKDRWAAIFK..........................................
SP-15/1-1140      KWSEDLDKYLHRFNELQNNLAPTN....................................
Carin5/1-1216     VWNEGINEFMKHYKNFEAWVNNYRALTSENVVTLESTKIDEGIINSWMEEENYERIGSVF
LIS04/1-1220      KWSESLDQYYHRFEELQNSLAPTT....................................

                                                                           330
RP13/1-1017       ........................................................KLDA
YerA41/1-1218     ........................................................DKNA
phiLo/1-1134      ............................................................
RHph_Y65/1-1067   ............................................................
EP_H11/1-1049     ............................................................
SP-15/1-1140      ............................................................
Carin5/1-1216     DKDLDSLDPETFEVFIKDVGEKNDPKNALVIREYVTGANLIDDPMLFMTEFFAQLKNENA
LIS04/1-1220      ............................................................

                            340
RP13/1-1017       DNIINLKDLC..................................................
YerA41/1-1218     YEITSIKDLL..................................................
phiLo/1-1134      ..........................EYLL..............................
RHph_Y65/1-1067   EKSPKLYNACRQANMEAI.........YLL..............................
EP_H11/1-1049     ....DMKKRCTPEQIAEIQSGEFNIEHHLA..............................
SP-15/1-1140      ...................AGSLREEHKVL..............................
Carin5/1-1216     PE..........................................................
LIS04/1-1220      ...................AGNLRNEHKVLKEVKSIEKFAEYMFSLLVESINAKVEEDKI

RP13/1-1017       ............................................................
YerA41/1-1218     ............................................................
phiLo/1-1134      ..................................VSPHSLKQTKIK..............
RHph_Y65/1-1067   ..................................KEDTAEKITGGKKNPKVTRYRTKQAD
EP_H11/1-1049     ..................................LKDEYEKLV..EAAPTKAAKNRVPVN
SP-15/1-1140      ..................IQEKSVKALADYMILQVEESLNRPTKEGHAKTKDIPGTLELI
Carin5/1-1216     ............................................................
LIS04/1-1220      SSHDSVQLIELLKTNLMD..........................................

                                  350
RP13/1-1017       .......RI..GDLNGLYSY......................GS................
YerA41/1-1218     .......SS..DDKSTLIKLLGLTETDKVKDRSRLEFKLTDDGDKKEPTYKLTTYVVETY
phiLo/1-1134      .......KI..ESLL.............................................
RHph_Y65/1-1067   HADYISAKI..SALR.............................................
EP_H11/1-1049     FNQFMLDKI..AELS.............................................
SP-15/1-1140      RSTLSNDNIAWSDMK.............................................
Carin5/1-1216     ............................................................
LIS04/1-1220      ......DSISWSDTK.............................................
```

19

```
                           360                               370
RP13/1-1017       .....PAIKGILDWMLE..GYS......................REEV............
YerA41/1-1218     FDLLLDPIKFVEKAMEH..SQKYEQSFFKFFMTYQDRLEGKFKLTNLL............
phiLo/1-1134      ................EKLGFD......................VEEA............
RHph_Y65/1-1067   ................E..FCS......................DEDI............
EP_H11/1-1049     ................E..ELE......................ESEI............
SP-15/1-1140      ................E..VME......................NFKVFDNRQRNVAIAA
Carin5/1-1216     ..........................FFKFFSTYEDRIEGKFVISNLM............
LIS04/1-1220      ................E..VLE......................NYEIFDKRKRNCLIAS

                           380
RP13/1-1017       ....QEVITHYP.............Y......................E.......WGGI
YerA41/1-1218     ....TYFIKNY..............N......................E.......WQSC
phiLo/1-1134      ....KRIFSFVL.............S......................QPGNPLIGWATV
RHph_Y65/1-1067   ....QRGFTKFP.............Y......................E.......WAAV
EP_H11/1-1049     ....IYLASFYP.............Y......................E.......WGAI
SP-15/1-1140      WKMVKLIERF...............YTTEEEFNLIDGLVAKAINDRLEKDDYSFVSYGLV
Carin5/1-1216     ....KYYQGGY..............N......................E.......WQAC
LIS04/1-1220      WKMFRLIERYYPKYEDDGTEYEKLDY...........LVAKAILDRIDEGDFGFISYGLI

                        EXO III
                   390       400       410                   420       430
RP13/1-1017       PKQLLGIYCAYDAGYTVLLANKMY...IPEME.F..........AYQVYIRHPYLATKFE
YerA41/1-1218     GIGDMGYYCILDGYYTVKLKEMLYDTNYDGCA.D..........GYIYYLYQSHFASLLE
phiLo/1-1134      PKEVLARYNAEDVRWTHEL.YLFY...TSLYPWE..........VWEVYQKQEELGSLMH
RHph_Y65/1-1067   PPKILGEYCCYDSYYTLGLKNHLW....DKYK.S..........VYQYYITQGWLGSTME
EP_H11/1-1049     PREILGEYCCYDSAYGVFI.HEMF...YDKYK.D..........GYNVYMLHPYLASIFE
SP-15/1-1140      PMKIIAPYGALDAVATVDL.KKYY...YKRMKEESETLGLDLLQGYSNWMKHFKMAYVME
Carin5/1-1216     GIGDMGYYCILDSYYTLKLKEMLYDTNHQGVK.N..........GYPFYVSQSNLAGLIE
LIS04/1-1220      PLRIIAPYGALDAVACIDL.RNYY...NERMTVESKELEIDLFKGYSNWMKHFKMAYIME

                            440       450       460          470       480
RP13/1-1017       VNGILWD.....DKRATDLETDTYTIMLKLLHDIIQDCDI...PLEKKLEATDILNRP.L
YerA41/1-1218     ANKVHIDKSRFVDWMKWAYETRMYHAKKVALHPKVVENYS...KSLYKKKFSGVIKTN.I
phiLo/1-1134      QVRIRYD.....VEAAKRLRGEFSRIAVESFREMISSPRLTKEAIEGRLRNLKVRDVPEL
RHph_Y65/1-1067   SYGLNWD.....DDKASEHERFYLVEATTCLNQIIQKLDFGEKTLDIHTRASMIFNDPSL
EP_H11/1-1049     ANGIRWD.....DEEATKEYDYCMSKMSEHLVSLIPNLKL...PPDQMLTARDFIYRE.L
SP-15/1-1140      RNGAFWN.....EELVKKERKFLEGTAVEATRTLHKSPLMADFLAEKKSGSFATFVVTKM
Carin5/1-1216     AYKVSINYDRFSKWKEWNLTERFRLAKEIALYPAVIDNYI...EGHYLKQKSTAYNKA.I
LIS04/1-1220      RNGAYWQ.....EDLVKKEKEFLEGAAIDATRALHTSPLMADYLVEKKAGDFSTYIVTQA

                                                              490
RP13/1-1017       P........................................YTVTWYTEKTRA...ERQ.
YerA41/1-1218     E........................................SVIKSYNGNKRVVAAELS.
phiLo/1-1134      KEALSLAYRVHLRAYKEATK..........DESIVEKYKKAYQKLLSSP...........
RHph_Y65/1-1067   P........................................D..................
EP_H11/1-1049     P........................................YENVWYTPGGQ....ERR.
SP-15/1-1140      PDAISRSREIFISDIRTKKRSKKKKEVVFSDGSVIEMTSTAILDLLTPEELEN...AKQA
Carin5/1-1216     E........................................SLILGYTKKKRVVARDIK.
LIS04/1-1220      PQAISESCQLFISDIIKKPRSKNKKQVVFSNGDTYDMTSPAISNLLTPEIREK...VKSD

                      500                                         510
RP13/1-1017       ......YVV..................................ATVGQKIEELK......
YerA41/1-1218     ......FVIKYQGKDLEEMIKTYYEHLAVGRKALVDHINETGNAKL.TKTQKGK......
phiLo/1-1134      ........V..................................LSPESEPADLSHLVVSK
RHph_Y65/1-1067   ..............................................ARKLIDLK......
EP_H11/1-1049     ......KMV..................................TTLAGKIADLS......
SP-15/1-1140      WIKDMNDTI..................................FSEFTKISEFK......
Carin5/1-1216     ......FV....................................................
LIS04/1-1220      WIKYLADNV..................................FSKFTKIKEFK......

                                                          520       530
RP13/1-1017       ....................................GIFNPGSNTDATRDKFWNPY....
YerA41/1-1218     ....................................IIGKEFTNDDKIIHKFIEIIDKKY
phiLo/1-1134      RESLLNSLRERLREEFERDLSMYGKENGEAWVSRMAAWFNPSSS.........KAY....
RHph_Y65/1-1067   ....................................GIFNPGTPDKLKQKPFWDKY....
EP_H11/1-1049     ....................................GFFNPNSNDKANKEKFWKAY....
SP-15/1-1140      ....................................EYFNPGSP..ANKTYLNKIL....
Carin5/1-1216     .......................................REF..................
LIS04/1-1220      ....................................DYFNPGSP..AVKEFLGKHL....
```

20

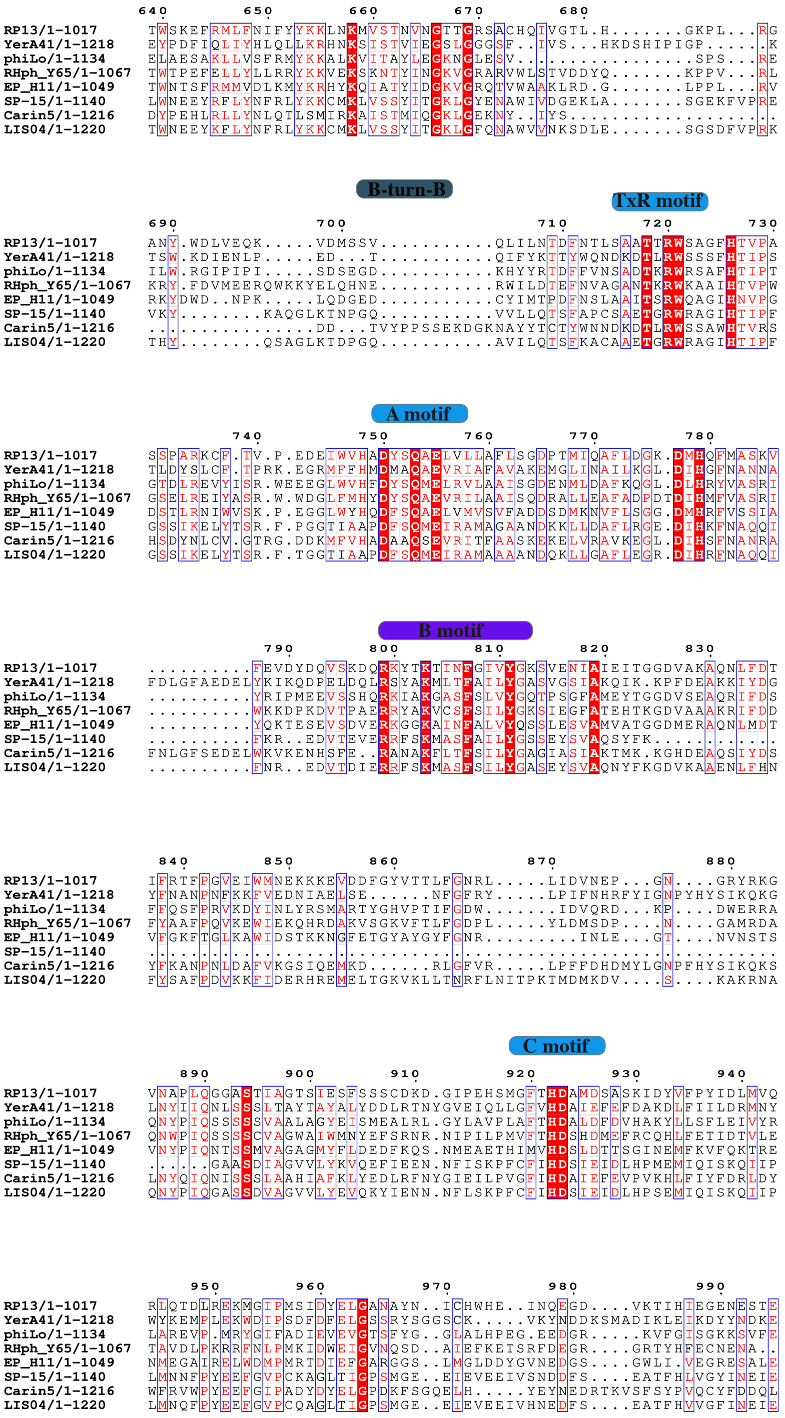
B-turn-B
TxR motif
A motif
B motif
C motif

```
                                  540       550
RP13/1-1017         .LT.........DEIQLGTVLLLFLEDIEL..............................
YerA41/1-1218       KLAFFGTNPPIADRIAVISQFINHLYNRAEGMV...........................
phiLo/1-1134        .VS.........VFHKAIS.....CMPLIA..............................
RHph_Y65/1-1067     .VT.........DFTTAISIYLVLSEQILM..............................
EP_H11/1-1049       .LT.........DEVEMCTMLWSVMDQLKVHDY...........................
SP-15/1-1140        .VN.........DEVRAARFLSKVKNELFTQDN...........................
Carin5/1-1216       ...........................NAENFVKEYFDFTELFKSNLEELKEYKARSTKK
LIS04/1-1220        .VN.........DEVRAAKFLFELKNKVFTIEN...........................

RP13/1-1017         ............................................................
YerA41/1-1218       ............................................................
phiLo/1-1134        ............................................................
RHph_Y65/1-1067     ............................................................
EP_H11/1-1049       ....................................WD......................
SP-15/1-1140        ....................................FSVDNYPASDRMFLKAVQSILDRM
Carin5/1-1216       VGKRESDKILMAGMNDKQKSIIAIQDLVNKKIQLGVIENSDYNGKLKAITSSVSMLQSKA
LIS04/1-1220        ....................................YSPDAYPDSNRVFLLAVERILDSM

                               560               570
RP13/1-1017         .......KGLLPDLYVA.......LGDKSF.IYQ.......................NKP
YerA41/1-1218       .......NITYEDVKQEYLDRVNSSTDYYD.IDRYLEFSSTRHEARLRMTNAVIPAGHEM
phiLo/1-1134        .......ANYIAEVLVK.......RVPK................................
RHph_Y65/1-1067     .......SPIIPDQAIE.......LVDR................................
EP_H11/1-1049       KTLLEISKGFVPKDSEK.......LEKEFF..............................
SP-15/1-1140        SGIKKAVADLEEDEIID.......NPEW................................
Carin5/1-1216       EAISTS.EVDWDALKQEHVDFINDEYCTYEKLDRRLDFGSTKDSTRLK............
LIS04/1-1220        KGIKKSVADLDEDEMID.......NPEW................................

                      580           590
RP13/1-1017         AVVMEAILGVTQ....QNNAL.................................GKGVQQ
YerA41/1-1218       VILQFLLVNRLKSTSIDHEFI.................................DKMPSD
phiLo/1-1134        ...........................................................E
RHph_Y65/1-1067     ...........................................................Q
EP_H11/1-1049       ............................................................
SP-15/1-1140        ...........................................................A
Carin5/1-1216       ...................FIKNGIMQGNMSILTLRKGLINRLVLSNNGLIDEVKEVPID
LIS04/1-1220        ...........................................................S

                   600
RP13/1-1017         SLN......FAV............................N...................
YerA41/1-1218       EFG......EYL............................V...................
phiLo/1-1134        DLSLLEEELGPI............................NTWPYKWEEVASKIFGEDWR
RHph_Y65/1-1067     DLNKAIENLMSV............................D...................
EP_H11/1-1049       .RE......FPV............................D...................
SP-15/1-1140        DLDEEEE..VTV............................D...................
Carin5/1-1216       DFM......KYLSAKMKEYAGSYEPDHSDFLKQLKASKVYT...................
LIS04/1-1220        DLDEAEE..VVV............................N...................

                               610
RP13/1-1017         .TLPE........N..I.....................................RR...F
YerA41/1-1218       .DLVKRGNLAEAGQ..G.....................................KR...L
phiLo/1-1134        ARIYP........T..MLQMWESIASG................................L
RHph_Y65/1-1067     .WTKY........D..GV...DLKTAKSIQKLM.........MDLIGTLDEKIGWKLYAF
EP_H11/1-1049       .EILT........R..IIQFGDTAESKHVRELIAMCV.....QDGLGAYEERFTNK...F
SP-15/1-1140        .KITA........KEGFAEYIEVLRTARLES...........PDLVNMFAECLNWDLPGT
Carin5/1-1216       .D..........................................................
LIS04/1-1220        .KIST........REGFLQYLELLKNSRLQSTGTIKSGGIELPDLVEVFAECLNYNLTST

                         620         630
RP13/1-1017         A..GPVVKFQYKMHNTF..L.......KL.................NPDDE........S
YerA41/1-1218       SAADKIILLEVKTAKGT..LSKILSGEKLGKHALRTIKQAWDFKNGRKNSL........D
phiLo/1-1134        N..DEYMEYLYSLFKDY..L.......GE.................DLNDYPACLERVPK
RHph_Y65/1-1067     R..SDVLEWQYKAHETY..G.......GL.................DIDNR........E
EP_H11/1-1049       K..ADVLDAQLAIHQLM..L.......GV.................NPDDE........S
SP-15/1-1140        K..EMDM...IRLHETYQLM.......GI.................DIEDE........S
Carin5/1-1216       .....................RLIQGNKFGKMPFRFIKESYEDAYGEGKSI........D
LIS04/1-1220        K..EPDI...IKLHEIYQLM.......GI.................DIEDE........S
```

22

```
RP13/996-1059         ..............................LLIKKLDKSTTYKVLSSEVIKSKSEAT...
YerA41/1199-1257      ..........................DILKLMKESFNIIEDSLKKEDHPVEKHPFVFMID
phiLo/1116-1175       .............RVLKRLEKFYEVEVLE...LSEGEETI...PLADIFAPKKALS...T
RHph_Y65/1047-1109    ..............MMPVFDRLKNYFDIEYEVTSRETERA...SLKEMFVARRAFS...K
EP_H11/1029-1095      .............KLLEALDQYSQWTPYV.EILKEKEKLN...SIANSFNSKSGIRSDGT
SP-15/1114-1177       HLADVWKVAYPKVVMEDVYDDAGN.PKIEDQYISRH..........ELFFPKRAFS...R
Carin5/1212-1271      .................DTVLRRFRNAFKISNEKVERLDDFVE.GKFDYIFKSAEPYN..
LIS04/1194-1257       HLAEVWKRAYPQVSMEDVYKDNGE.LRIEDQYIERS..........ELFFPKRAYS...R


RP13/996-1059         SWAEMFTTGTALK...DSWGKTVTKNTVELKVQMLPQTCM
YerA41/1199-1257      GVKPDC...RFREEVTDQYVYSAKIGLK............
phiLo/1116-1175       LIGKTVP..TVSGKIALREVSDES................
RHph_Y65/1047-1109    YLGTDVT..VVGGNLKLVKKNGIKI...............
EP_H11/1029-1095      GWGCKYI..ELKAKITLEPKHGEKAA..............
SP-15/1114-1177       YTGT..T..RKFGH....KKIHITIK..............
Carin5/1212-1271      .MNTYRSIPQYKLSGEIEVTW...................
LIS04/1194-1257       YTGT..T..RQFGH....KKIHITTR..............
```

**Supplementary Fig. 1**

Multiple sequence alignment (MSA) of A-family DNA polymerase members fused to an active uracil DNA glycosylase (UDG) domain, the SP15-like DNAPs. Sequences were identified using BLASTp against DNAP from YerA41 with the BLOSUM 40 matrix. After selecting sequences that covered at least 80% of the query, alignment was performed using MAFFT web server[1]. The final MSA was visualized using ESPRIPT[2]. Sequences with a UDG-B motif containing "NP/MY" belong to TDG/MUG family, while sequences containing "HP" belong to type I / type III / type IV / type V family. Abbreviations. RP13: *Ralstonia* phage 13, YerA41: *Yersinia ruckeri* phage 41, phiLo: *Thermus phage phiLo*, Rhph_Y65: *Rhyzobium* phage Y65, EP-H11: *Escherichia* phage H11, SP-15: *Bacillus* phage 15, Carin5: *Cobetia marina* phage 5, LiS04: *Listeria* phage.

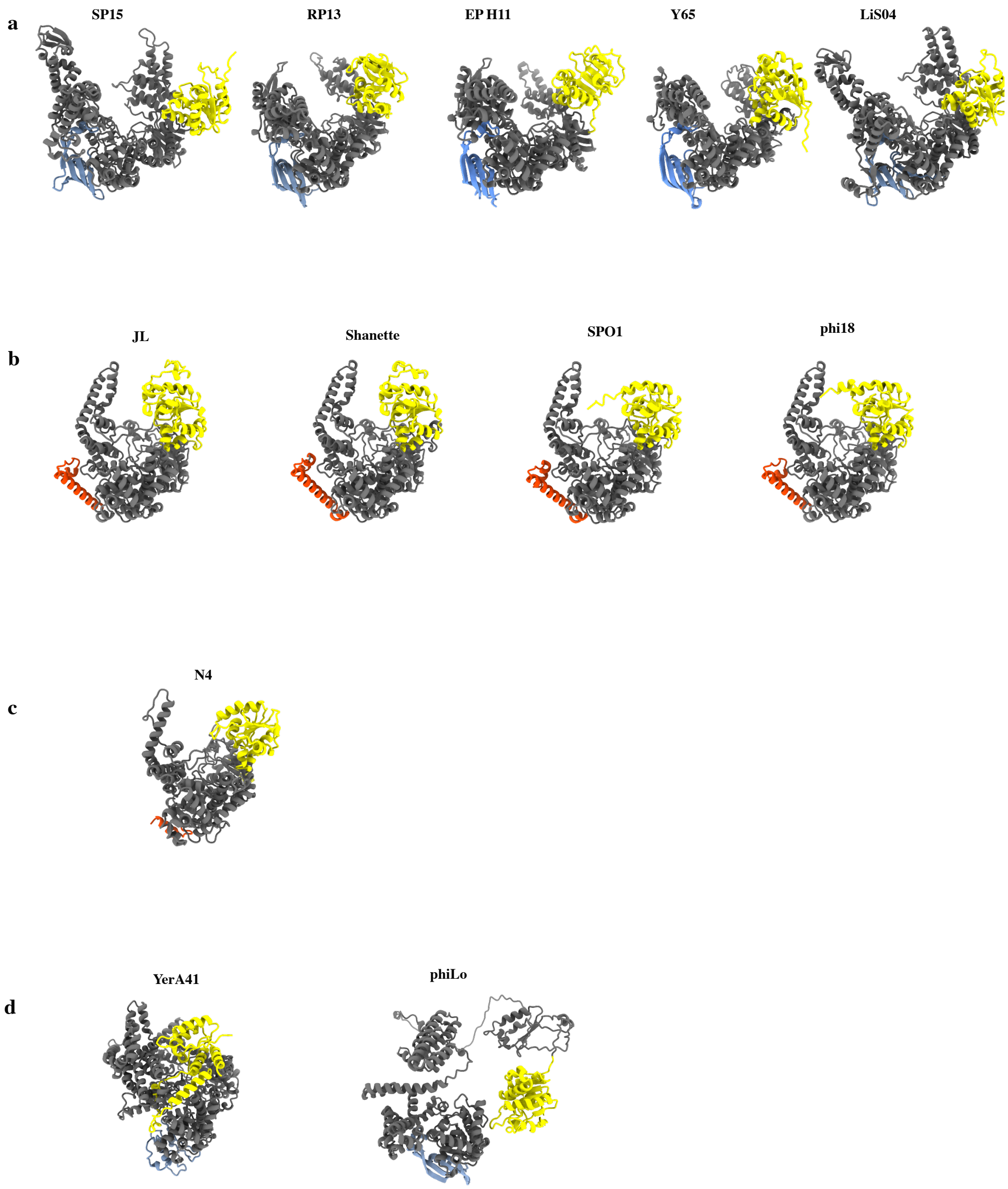


**Supplementary Fig. 2**

**a.** Alphafold3 models of different members of SP15-like DNAP sub-family. The CTD domain is colored in blue, the UDG domain is colored in yellow.

**b.** Alphafold3 models of different members of SPO1-like sub-family. The CTD helix is colored in red, the UDG domain is colored in yellow.

**c.** Alphafold3 model of N4 DNA polymerase. The CTD helix is colored in red, the UDG domain is colored in yellow. The UDG domain lacks the catalytic residues.

**d.** AlphaFold2 models of YerA41 and phiLo DNA polymerases. The UDG domains (yellow) are confidently modeled, as well as the canonical DNA polymerase cores (gray). However, insertions are not confidently modeled. Only the CTD domain (blue) of phiLo DNA polymerase is reliably predicted.

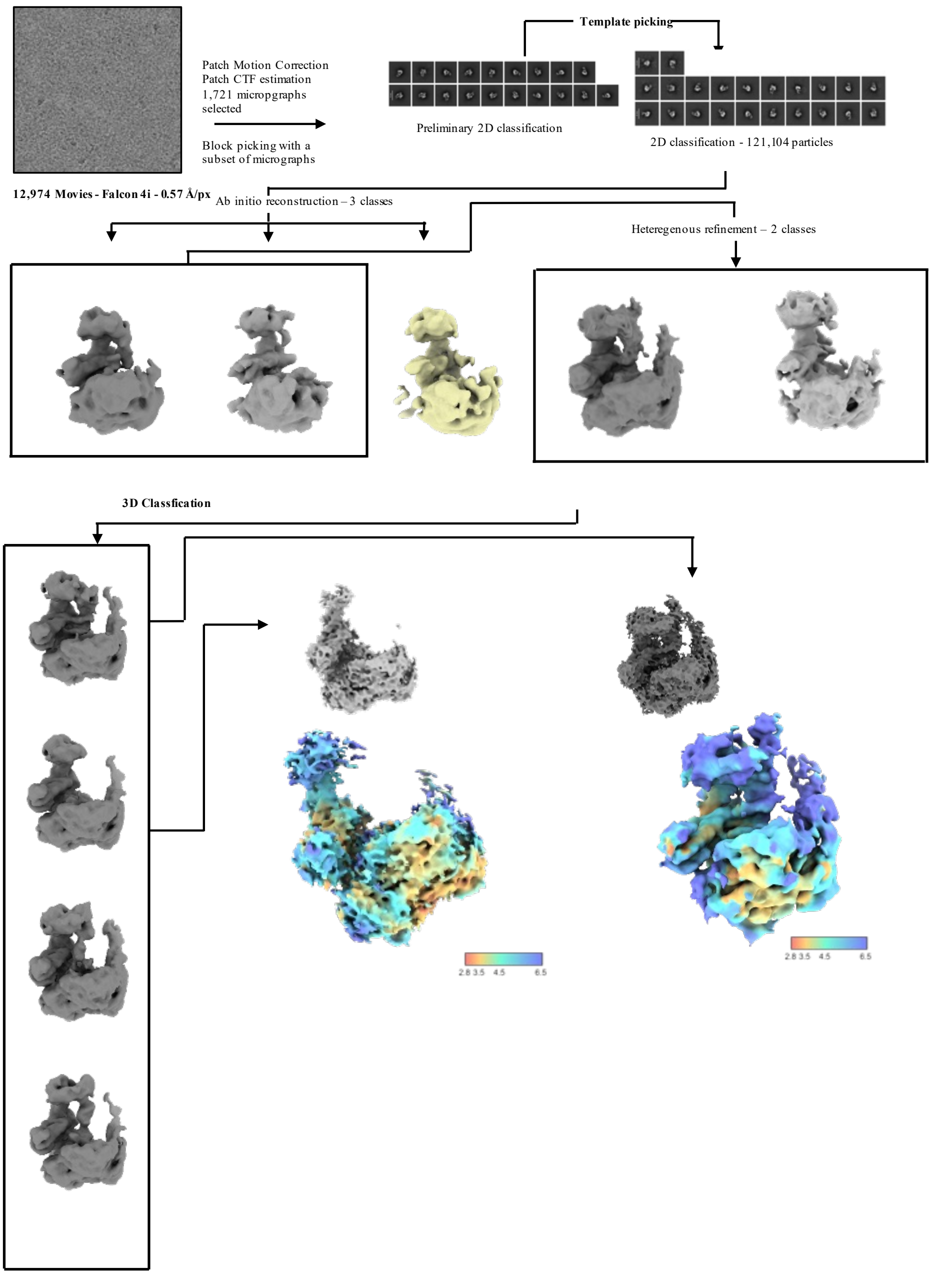


**Supplementary Fig. 3**

Workflow of cryo-EM data processing of DNA polymerase from *YerA41* DNAP with DNA substrate oSM72/oSM073.

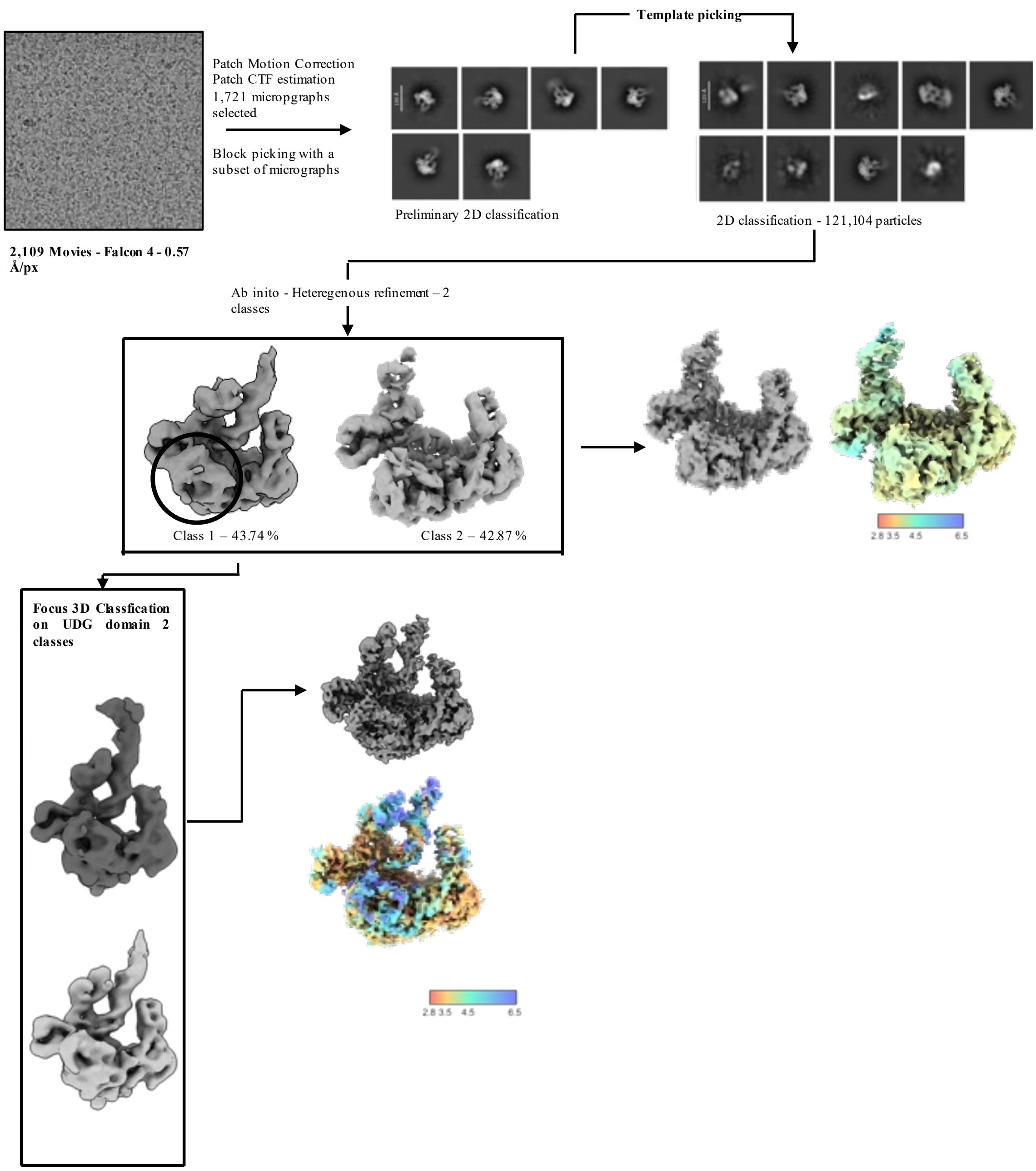


**Supplementary Fig. 4**

Workflow of cryo-EM data processing of DNA polymerase from *phiLo* DNAP with DNA substrate oSM72/oSM073.

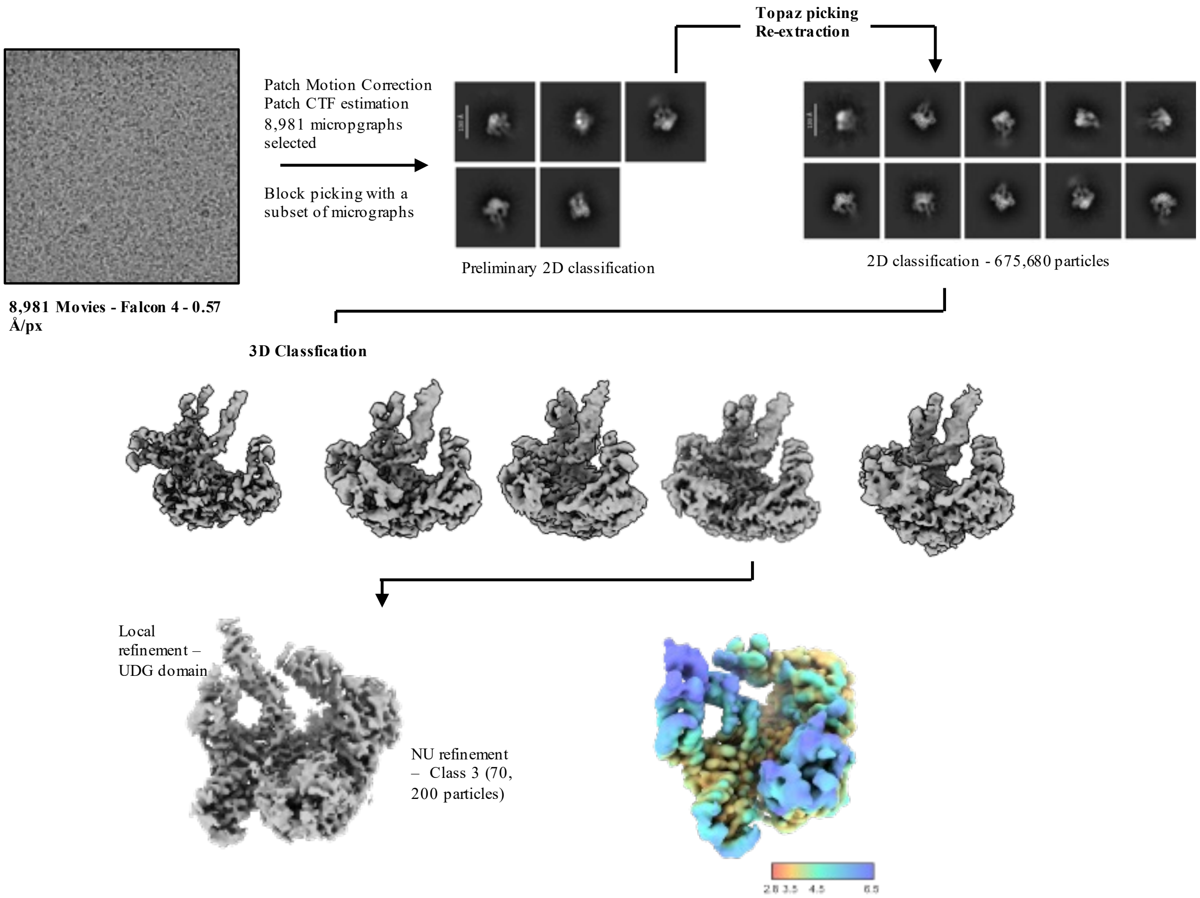


**Supplementary Fig. 5**

Workflow of cryo-EM data processing of DNA polymerase from *phiLo* DNAP with DNA substrate oSM72/oSM074.

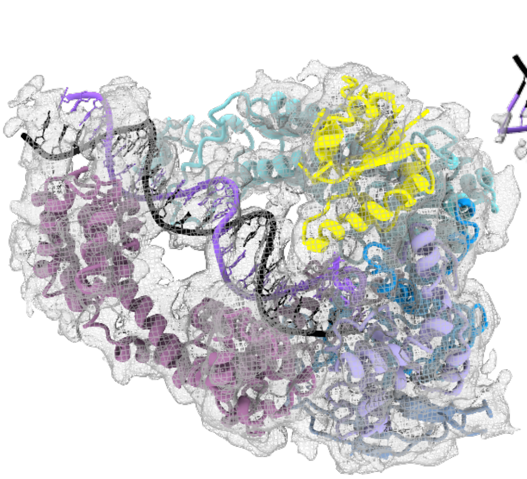

**Fit DNAP I YerA41**
**into cryo-EM map**

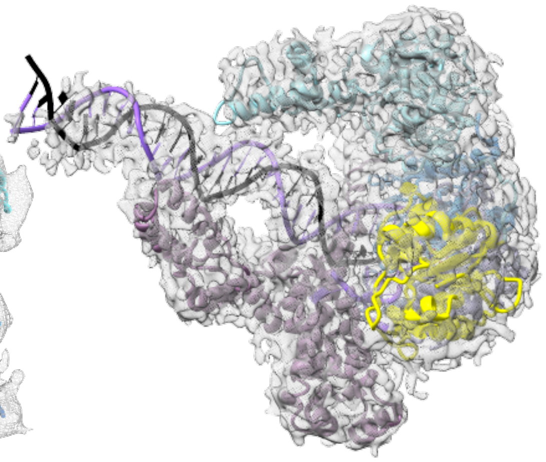

**Fit DNAP I phiLo**
**into cryo-EM map**

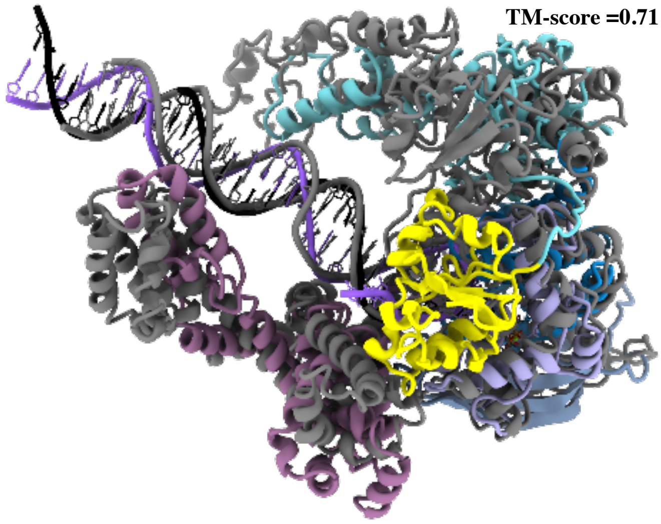


**Supplementary Fig. 6**

**a.** Fit of YerA41 DNAP ternary complex with unmodified template-primer DNA duplex in cryo-EM map.

**b.** Fit of phiLo DNAP ternary complex with unmodified template-primer DNA duplex in cryo-EM map.

**c.** Superimposition of DNAP from phiLo (color-coded as in **Fig. 1e**) and YerA41(in grey).

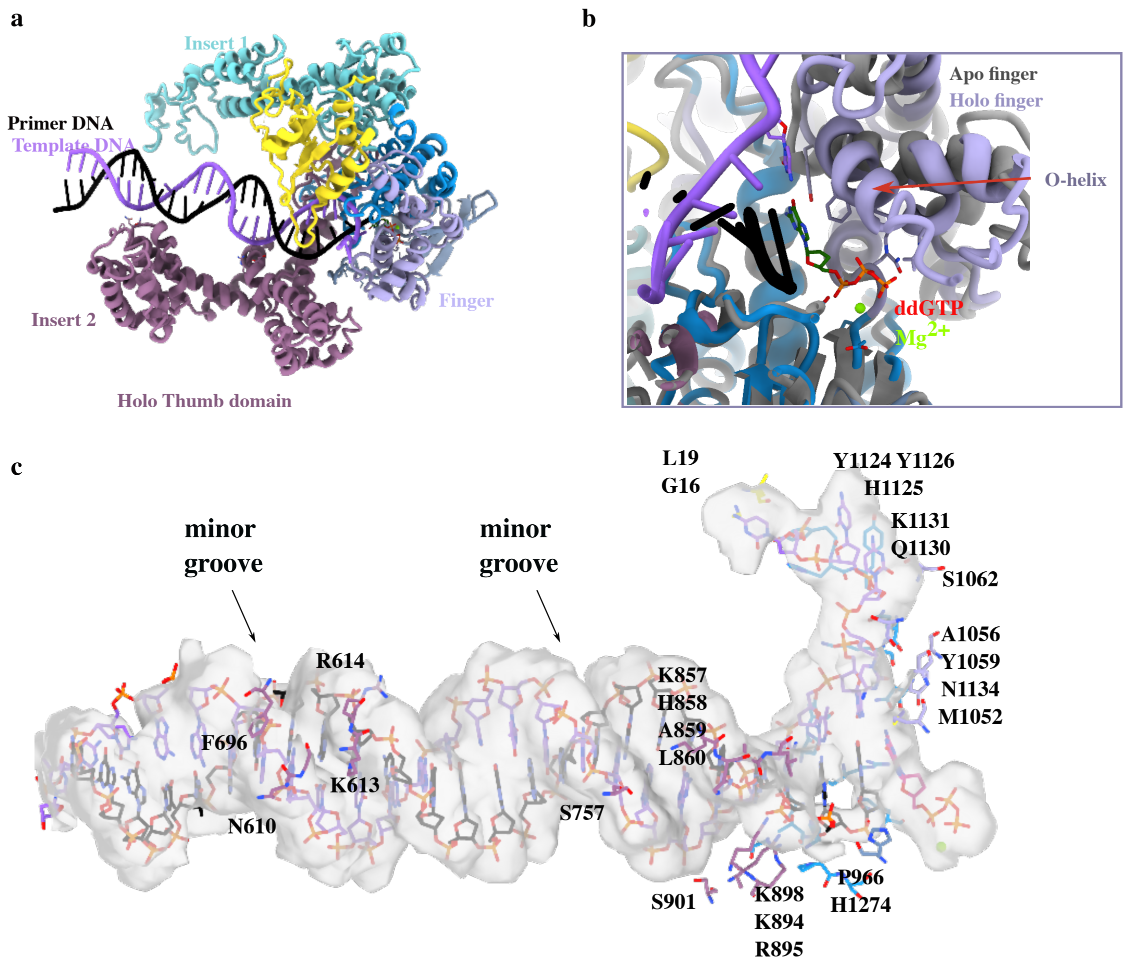


**Supplementary Fig. 7**

**a.** Rearrangement of Insert2 and Finger domains upon DNA binding and incoming nucleotide.

**b.** Movement of the O-helix from the Finger domain to stabilize the incoming nucleotide, capturing the elongation state of the enzyme. The structure of the ternary complex is colored, the apo conformation is in grey.

**c.** Experimental cryo-EM density represented in transparency surface focused on the DNA substrate stabilized by multiple contacts of YerA41 DNA polymerase (the residue numbers involved are in black). Nucleotides are represented in the stick mode residues and color-coded as in **Fig. 2.** Point contacts occur in majority in the minor groove of the DNA duplex, indicated with a black arrow.

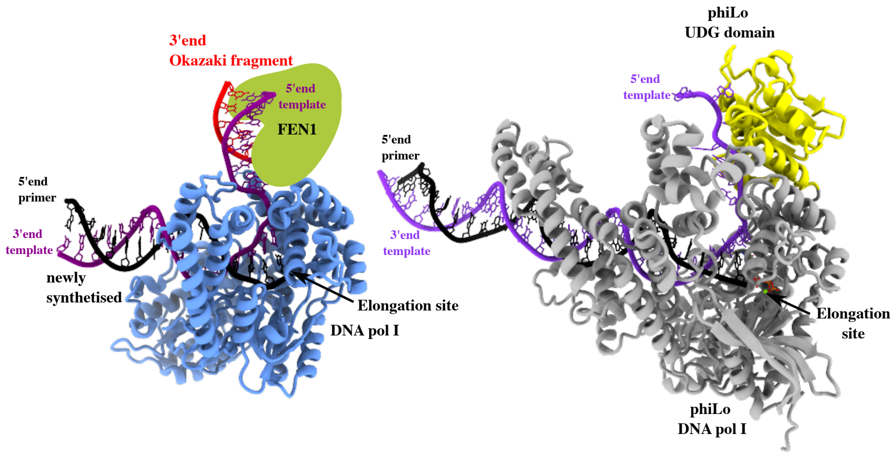


**Supplementary Fig. 8**

Assembly of *E. coli* DNA polymerase I with the endonuclease Fen1, as reported in Botto *et al.*,[3] in complex with a DNA duplex in elongation mode. The template strand is colored in purple, the newly synthesized primer is colored in black, and the Okazaki fragment, running from 5’ to 3’, is colored in red. The Fen1 protein is not present in the PDB structure (PDB ID 8OO6), but its position can be modeled using the position of the Okazaki fragment ahead of the elongation site. This assembly is compared with the structure of phiLo DNAP I fused to its UDG domain (colored in yellow) in complex with the DNA duplex. In this structure, the template strand is colored in purple, and the primer strand is colored in black. The localization of the UDG domain is the same as the modelled position of FEN1.

| **Data collection and processing** | #1 phiLo Elongation mode<br>with dU-containing template<br>(EMD-52894)<br>(PDB 9QHS) | #2 phiLo<br>Apo mode<br><br>(EMDB-53178)<br>(PDB 9QHT) | #3 phiLo<br>Elongation mode<br>with dsDNA<br><br>(EMDB-53179)<br>(PDB 9QHU) | #4 YerA<br>Elongation mode<br>with dsDNA<br><br>(EMDB-53227)<br>(PDB 9QLH) | #5 YerA<br>Apo mode<br><br>(EMDB-53228)<br>(PDB 9QLI) |
|---|---|---|---|---|---|
| Magnification | 265,000 | 265,000 | 265,000 | 215,000 | 215,000 |
| Voltage (kV) | 200 | 200 | 200 | 300 | 300 |
| Electron exposure (e–/Å$^2$) | 40 | 40 | 40 | 40 | 40 |
| Defocus range (μm) | -1 to -3 | -1 to -3 | -1 to -3 | -0.8 to -2 | -0.8 to -2 |
| Pixel size (Å) | 0.56 | 0.56 | 0.56 | 0.57 | 0.57 |
| Symmetry imposed | C1 | C1 | C1 | C1 | C1 |
| Images (no.)<br>Initial particle images (no.) | 8,981<br>675,680 | 2,109<br>121,104 | 2,109<br>121,104 | 12,974<br>822,203 | 12,974<br>822,203 |
| Final particle images (no.) | 70,200 | 34,104 | 41,506 | 30,217 | 51,255 |
| Map resolution (Å)<br>FSC threshold | 2.99<br>0.143 | 3.6<br>0.143 | 3.57<br>0.143 | 4.86<br>0.143 | 3.66<br>0.143 |
| Map resolution range (Å) | 1.49-5.02 | 2.5-5.5 | 2.76-4.25 | 3.69-7 | 3.28-6 |
| Map sharpening *B* factor (Å$^2$) | 130 | 150 | 115 | 121 | 120 |
| **Refinement** | | | | | |
| Model resol. (Å)<br>FSC threshold | 3.1<br>0.143 | 3.6<br>0.143 | 3.8<br>0.143 | 4.1<br>0.143 | 4.5<br>0.143 |
| Model resolution range (Å) | 2.6-3.9 | 3.5-3.9 | 3.8-4.2 | 3.9-7.6 | 3.8-8.1 |
| Model composition<br>Non-H atoms<br>Protein residues<br>Nucleotides<br>Ligands | <br>10592<br>1140<br>66<br>$Mg^{2+}$ ddCTP | <br>9048<br>1114<br>0<br>0 | <br>10363<br>1111<br>64<br>$Mg^{2+}$ ddGTP | <br>11525<br>1257<br>56<br>$Mg^{+2}$ ddGTP | <br>10361<br>1257<br>0<br>0 |
| *B* factors (Å$^2$)<br>Protein<br>Nucleotides<br>Ligand | <br>42.04<br>53.08<br>67.35 | <br>44.2<br>-<br>- | <br>81.95<br>105.54<br>116.02 | <br>109.4<br>193.9<br>260.4 | <br>130.0<br>-<br>- |
| R.m.s. deviations<br>Bond lengths (Å)<br>Bond angles (°) | <br>0.01<br>1.224 | <br>0.008<br>1.49 | <br>0.005<br>1.161 | <br>0.011<br>1.441 | <br>0.007<br>1.217 |
| Validation<br>MolProbity score | <br>2.70 | <br>2.28 | <br>2.90 | <br>3.23 | <br>2.77 |
| Ramachandran plot<br>Favored (%)<br>Allowed (%)<br>Disallowed (%) | <br>90.51<br>8.96<br>0.53 | <br>92.16<br>7.48<br>0.36 | <br>88.60<br>10.68<br>0.72 | <br>92.66<br>7.18<br>0.16 | <br>91.37<br>7.35<br>1.28 |
| Model vs Data<br>CC (box)<br>CC (main chain)<br>CC (side chain)<br>CC (ligand) | <br>0.65<br>0.59<br>0.60<br>0.53 | <br>0.72<br>0.67<br>0.66<br>- | <br>0.75<br>0.66<br>0.65<br>0.77 | <br>0.71<br>0.56<br>0.59<br>0.69 | <br>0.55<br>0.49<br>0.51<br>- |

**Supplementary Table 1**: Statistics of Cryo-EM data collection, model refinement and validation.

| molecule | locus | start | end |
|---|---|---|---|
| Enterobacteria phage T7 NC_001604.1 | T7p29 NP_041982.1 | 14353 | 16467 |
| Enterobacteria phage T7 NC_001604.1 | T7p10 NP_041963.1 | 6449 | 7588 |
| Bacillus phage Shanette NC_028983.1 | SHANETTE_176 YP_009216171.1 | 2779 | 5557 |
| Bacillus phage Shanette NC_028983.1 | SHANETTE_164 YP_009216159.1 | 109710 | 112544 |
| Thermus phage phiLo MH673673.1 | phiLo_118 AYJ73970.1 | 122999 | 124138 |
| Thermus phage phiLo MH673673.1 | phiLo_141 | 150661 | 151695 |
| Thermus phage phiLo MH673673.1 | phiLo_158 AYJ74010.1 | 166090 | 169617 |
| Rhizobium phage RHph_TM30 NC_070966.1 | EVB93_314 YP_010671079.1 | 142768 | 144666 |
| Rhizobium phage RHph_TM30 NC_070966.1 | EVB93_322 YP_010671087.1 | 149854 | 153183 |
| Rhizobium phage RHph_TM30 NC_070966.1 | EVB93_050 YP_010670835.1 | 22406 | 23362 |
| Rhizobium phage RHph_Y65 NC_070967.1 | EVB97_313 YP_010671442.1 | 141944 | 143842 |
| Rhizobium phage RHph_Y65 NC_070967.1 | EVB97_321 YP_010671450.1 | 149020 | 152349 |
| Rhizobium phage RHph_Y65 NC_070967.1 | EVB97_048 YP_010671197.1 | 21647 | 22603 |
| Yersinia phage YerA41 MW570730.1 | YerA41_127 QSM00828.1 | 68280 | 70193 |
| Yersinia phage YerA41 MW570730.1 | YerA41_144 QSM00845.1 | 85132 | 89052 |
| Yersinia phage YerA41 MW570730.1 | YerA41_158 QSM00859.1 | 99157 | 100053 |
| Bacillus phage SPO1 NC_011421.1 | SPO1_146 YP_002300417.1 | 102241 | 105900 |
| Bacillus phage SPO1 NC_011421.1 | SPO1_157 YP_002300428.1 | 110411 | 111682 |
| Listeria phage LIS04 OQ999172.1 | LIS04_113 WJZ23541.1 | 106544 | 110317 |
| Listeria phage LIS04 OQ999172.1 | LIS04_116 WJZ23544.1 | 111851 | 112753 |
| Listeria phage LIS04 OQ999172.1 | LIS04_132 WJZ23560.1 | 124062 | 125951 |
| Bacillus phage SP_10 NC_019487.1 | SP10_191 YP_007003448.1 | 116989 | 119646 |
| Bacillus phage SP_10 NC_019487.1 | SP10_198 YP_007003455.1 | 34391 | 35899 |
| Ralstonia phage RP13 LC554890.1 | RP_ORF24 BCG50042.1 | 11963 | 12877 |
| Ralstonia phage RP13 LC554890.1 | RP_ORF13 BCG50031.1 | 5065 | 6927 |
| Ralstonia phage RP13 LC554890.1 | RP_ORF88 BCG50106.1 | 62706 | 65885 |
| Cobetia phage Carin5 PV155638.1 | CLPJFFLI_00164 XPT09303.1 | 100312 | 101181 |
| Cobetia phage Carin5 PV155638.1 | CLPJFFLI_00196 XPT09335.1 | 132883 | 134757 |
| Cobetia phage Carin5 PV155638.1 | CLPJFFLI_00181 XPT09320.1 | 115291 | 119106 |
| Escherichia phage EP_H11 OP688485.1 | WBU87675.1 | 37287 | 38045 |
| Escherichia phage EP_H11 OP688485.1 | WBU87676.1 | 38110 | 40053 |
| Escherichia phage EP_H11 OP688485.1 | WBU87645.1 | 18921 | 22208 |
| N4 phage | EPNV4_gp39 | 22821 | 25400 |
| Bacillus phage SP-15 NC_031245.1 | SP15_143 YP_009302530.1 | 126589 | 128448 |
| Bacillus phage SP-15 NC_031245.1 | SP15-298 YP_009302695.1 | 217771 | 218529 |
| Bacillus phage SP-15 NC_031245.1 | SP15_109 YP_009302497.1 | 102353 | 107194 |
| Herelleviridae sp. Phage BK042982.1 | DAT11781.1 | 15105 | 18248 |

| | | | |
|---|---|---|---|
| Herelleviridae sp. Phage BK042982.1 | DAT11811.1 | 26335 | 27012 |
| Herelleviridae sp. Phage BK042982.1 | DAT11782.1 | 37232 | 39022 |
| Bacteriophage sp. OP075190.1 | UWG86726.1 | 110 | 3649 |
| Bacteriophage sp. OP075190.1 | UWG87231.1 | 384705 | 385892 |
| Bacteriophage sp. OP075190.1 | UWG87052.1 | 245450 | 246973 |
| Escherichia phage vB_EcoM_IME392 MH719082.1 | AXY86062.1 | 7051 | 8994 |
| Escherichia phage vB_EcoM_IME392 MH719082.1 | AXY86063.1 | 9059 | 9817 |
| Escherichia phage vB_EcoM_IME392 MH719082.1 | AXY86073.1 | 24922 | 28209 |
| Caudoviricetes sp. BK045098.1 | DAP65209.1 | 7377 | 8150 |
| Caudoviricetes sp. BK045098.1 | DAP65173.1 | 138889 | 140703 |
| Caudoviricetes sp. BK045098.1 | DAP65174.1 | 145366 | 148965 |

**Supplementary Table 2**

List of genes for A-family DNA Polymerase, DNA Polymerase X, and DNA Ligase in each studied genome

This table provides a list of genes encoding A-family DNA polymerase, DNA polymerase X, and DNA ligase identified in the genomes analyzed in this study. Each entry includes the genome identifier, gene identifier, and relevant annotations.

| Name of oligonucleotides | Sequences | Function |
|---|---|---|
| iSM0051 | aacgcAAAGCCTTTATGTTCATCCG | primer YerA_exo- |
| iSM0056 | tgcgACCCGCAATTTTCCGTT | primer YerA_exo- |
| iSM0054 | tgcgACCAGCAAAGGTGAAACC | primer phiLo_exo- |
| iSM0055 | accgcGAAGCTCACCTCGTCCAG | primer phiLo_exo- |
| oSM0072 | CACAGACGTAGCAGCTGGAACCTCCGTA | Primer complex cryoEm |
| oSM0073 | AAAA[U]CCCCGCTACGGAGGTTCCAGCTG CTACGTCTGTG | Template dU complex cryoEM |
| oSM0074 | AAAATCCCCCCTACGGAGGTTCCAGCTGCT ACGTCTGTG | Template complex cryoEM |
| iSM0094 | FAM CCG CGT ATA GCC GGG GTT CCC GTA CCG | Primer activity |
| iSM0093 | AGGAGAGGGAGT[Spd]CGGTACGGGAACCC CGGCTATACGCGG | Template THF activity |
| iSM0106 | FAM AGCGCATCAGCTGCAG | Primer activity |
| iSM0104 | CY5GGATCCCCGGGTUCAGACCTGCAGCTG ATGCGCT | Template dU activity |
| iSM033 | CY5CTG CAG CTG ATG CGC [U] GTA CGG ATC CCC GGG TAC | Template dU |
| iSM034 | GTACCC GGG GAT CCG TAC GGC GCA TCA GCT GCA G | Complementary DNA |
| iSM035 | CY5 CTG CAG CTG ATG CGC [Spd] GTA CGG ATC CCC GGG TAC | Template THF |
| iSM036 | CY5 CTG CAG CTG ATG CGC | Template gap |
| iSM037 | GTA CGG ATC CCC GGG TAC | Template 2 gap |

**Supplementary Table 3**

List and sequences of primers and oligonucleotides used for this study.

### References


1. Katoh, K., Rozewicki, J. & Yamada, K. D. MAFFT online service: multiple sequence alignment, interactive sequence choice and visualization. *Briefings in Bioinformatics* **20**, 1160–1166 (2019).
2. Gouet, P., Courcelle, E., Stuart, D. I. & Metoz, F. ESPript: analysis of multiple sequence alignments in PostScript. *Bioinformatics* **15**, 305–308 (1999).
3. Botto, M. M., Borsellini, A. & Lamers, M. H. A four-point molecular handover during Okazaki maturation. *Nat Struct Mol Biol* **30**, 1505–1515 (2023).
